# Retinoic acid exerts sexually dimorphic effects over muscle energy metabolism and function

**DOI:** 10.1101/2021.06.04.447134

**Authors:** Yaxin Zhao, Marta Vuckovic, Hong Sik Yoo, Nina Fox, Adrienne Rodriguez, Kyler McKessy, Joseph L. Napoli

## Abstract

The retinol dehydrogenase Rdh10 catalyzes the rate-limiting reaction that converts retinol into retinoic acid (RA), an autacoid that regulates energy balance and suppresses adiposity. Relative to WT, *Rdh10*+/− males experienced reduced fatty-acid oxidation, glucose intolerance and insulin resistance. Running endurance decreased 40%. *Rdh10*+/− females increased reliance on fatty acid oxidation and did not experience glucose intolerance nor insulin resistance. Running endurance improved 2.2-fold. Estrogen increased, revealed by a 40% increase in uterine weight. Because skeletal muscle energy use restricts adiposity and insulin resistance, we assessed the mixed fiber type gastrocnemius muscle (GM) to determine the effects of endogenous RA on muscle metabolism in vivo. RA in *Rdh10*+/− male GM decreased 38% relative to WT. TAG content increased 1.7-fold. *Glut1* mRNA and glucose decreased >30%. *Rdh10*+/− male GM had impaired electron transport chain activity, and a 60% reduction in fasting ATP. The share of oxidative fibers increased, as did expression of the myogenic transcription factors *Myog* and *Myf5*. Centralized nuclei increased 5-fold in fibers—indicating muscle malady or repair. In *Rdh10*+/− female GM, RA decreased only 17%, due to a 1.8-fold increase in the estrogen-induced retinol dehydrogenase, Dhrs9. *Rdh10*+/− female GM did not amass TAG, increase oxidative fibers, decrease *Glut1* mRNA or glucose, nor increase centralized nuclei. Expression of *Myog* and *Myf5* decreased. Electron transport chain activity increased, elevating fasting ATP >3-fold. Thus, small decreases in skeletal muscle RA affect whole body energy use, insulin resistance and adiposity, in part through estrogen-related sexual dimorphic effects on mitochondria function.

## Introduction

All-*trans*-retinoic acid (RA) regulates embryonic development, cell proliferation, differentiation and numerous functions of differentiated cells through complex genomic and non-genomic mechanisms (1–4). RA biogenesis stems from either retinol (vitamin A) or β-carotene (5–8). As many as eight retinol dehydrogenases and reductases, of the short-chain dehydrogenase/reductase gene family, catalyze conversion of retinol into retinal, and retinal into retinol (9). At least three retinal dehydrogenases, of the aldehyde dehydrogenase gene family, convert retinal generated from retinol or β-carotene, irreversibly into RA. Cells co-express multiple retinoid metabolic enzymes, but gene ablations reveal dissimilar phenotypes for each, consistent with singular contributions to RA (10, 11). For example, knocking out the dehydrogenase *Rdh1* increases adiposity in mice fed a low-fat diet, from decreased RA in brown adipocytes, impairing lipolysis and fatty acid oxidation (12). In contrast, the dehydrogenase *Dhrs9* seems crucial to the hair follicle cycle, and its decreased expression correlates with squamous cell carcinoma (13, 14). Ablation of the dehydrogenase *Rdh10* causes lethality between embryo days 10.5-14.5 from impaired nervous system and craniofacial development (15–17). During postnatal growth, Rdh10 generates RA to support spermatogenesis and organ repair, and to regulate energy use (18–20). Ablation of the retinal reductase *Dhrs3* increases embryonic RA amounts by ~40% and causes death late in gestation from defects in heart development (21, 22). Consistent with partially overlapping contributions to RA homeostasis, reducing expression of one reductase/dehydrogenase can provoke compensation by others. These observations prompt questions about the needs for and functions of so many enzymes dedicated to RA homeostasis.

Establishing precise physiological functions of endogenous RA cannot be attained by feeding vitamin A-deficient diets or pharmacological RA dosing, which prompts RA toxicity. The heterozygote *Rdh10*-null mouse (*Rdh10*+/−) experiences a limited decrease in tissue RA (≤ 25%), allowing reproducible assessment of endogenous RA functions in vivo (20). When fed a high-fat (western) diet, *Rdh10*+/− males gain adipose and suffer liver steatosis, whereas females form adipocytes in bone. To develop further insight into the function of Rdh10 and physiological RA actions, we expanded evaluation of the *Rdh10*+/− mouse to include whole-body energy metabolism and skeletal muscle function. Skeletal muscle accounts for about half of all energy used, producing not only work but heat to defend body temperature (23). Exercise, through muscles’ bountiful demand for energy, ameliorates obesity and insulin resistance (24, 25). Energy metabolism, obesity, metabolic syndrome and muscle function interweave to affect overall health.

RA induces skeletal muscle development through controlling expression of myogenic regulatory factors, such as Myf5, MyoD and Myog during embryogenesis and in primary myoblasts and cell lines (26–29). RA induces proliferation of cultured myoblasts and sustains their survival, suggesting a contribution to skeletal muscle regeneration (30, 31). Pharmacological dosing with RARγ agonists stimulates skeletal muscle repair and reduces extent of fatty fibrotic tissue lesions in a muscle injury model (32). Although it exerts crucial actions during muscle development and repair, little has been revealed about RA function on muscle performance in vivo.

Here we report sexually dimorphic differences in *Rdh10*+/− mice in the respiratory exchange ratio (RER), running endurance, glucose tolerance and insulin resistance, and in the muscle electron transport chain. Decreased Rdh10 in male gastrocnemius muscle (GM), a mixed-fiber type muscle, associates with TAG accumulation, an increase in centralized nuclei, reduced expression and activity of complex IV components of the electron transport chain, and decreased ATP. *Rdh10*+/− females have increased *Cox5a* expression and complex IV activity, and a higher concentration of resting ATP than WT. Estrogen increased in *Rdh10*+/− females, as did expression of the estrogen-induced retinol dehydrogenase Dhrs9 (33). RA decreased in *Rdh10*+/− female GM less than half the male decrease, likely due to compensation by Dhrs9. These results provide new insight into the physiological functions of Rdh10, sex differences in RA biosynthesis and actions related to energy use, insulin resistance and muscle function.

## Results

### *Rdh10*+/− differ by sex from WT in fuel use

*Rdh10*+/− males fed a high-fat diet endure glucose intolerance and insulin resistance, but females were not assessed (20). Here we found that *Rdh10*+/− females of the same age, and fed the same diet since weaning, did not experience glucose intolerance nor insulin resistance (Supporting Information Figure S1, S2). Further, RER showed that WT males relied more on fatty acid oxidation than WT females during normal activity (Figure 1A). Relative to WT, *Rdh10*+/− males decreased fatty acid oxidation and increased carbohydrate use during the 12-h dark (feeding) cycle, but did not differ from WT males during the light cycle. Relative to WT females, *Rdh10*+/− females decreased carbohydrate use and increased fat oxidation throughout 24 h.

**Figure 1.**
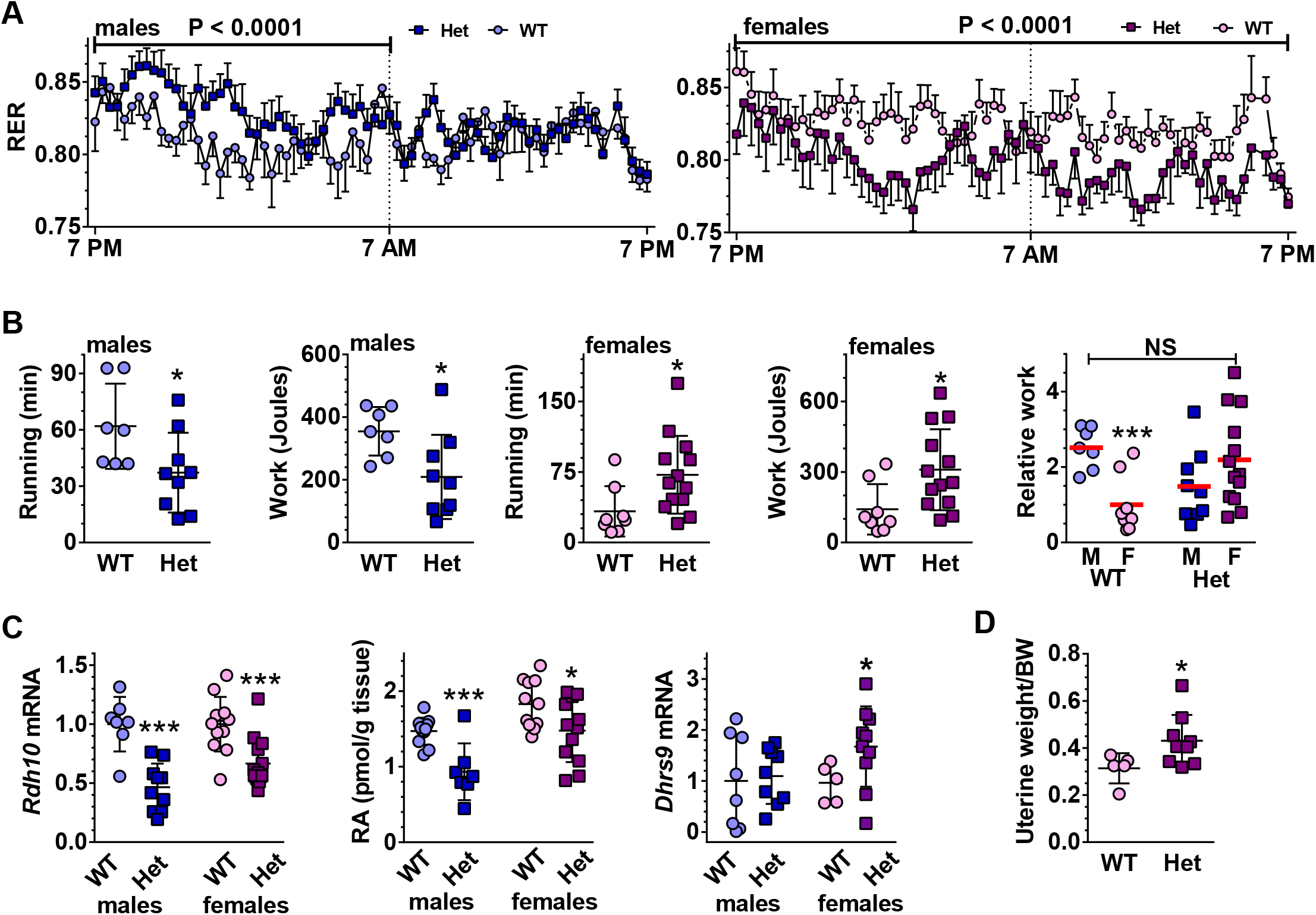
*Rdh10*+/− (Het) males and females differ from each other and WT in GM RA, fuel use, running endurance and myogenic transcription factor expression. *A*, Respiratory exchange ratio (CO_2_/O_2_) during ambient temperature and ad lib feeding. Data are means +/− SEM (n = 6 to 10). Data were analyzed by ordinary two-way ANOVA from 75 equally-spaced observations over 24 h for each sex/genotype. *B*, Running time and work done during the run-to-exhaustion test (n = 7 to 13). The last panel shows work done relative to female WT average of 1. Red bars denote averages. *C, Rdh10* mRNA (n = 7 to 14), RA content (n = 12), *Dhrs9* mRNA (n = 7 to 13) in GM. *D*, Weight of uteri relative to total body weight (BW) (n = 5 to 9). Symbols in all 7 figures: WT male, light blue circles; *Rdh10*+/− (Het) male, dark blue squares; WT female, light pink circles; *Rdh10*+/− (Het) female, dark purple squares.

### *Rdh10*+/− males and females differ in running endurance

Greater reliance on carbohydrates vs fatty acid oxidation implied possible endurance differences. In run-to-exhaustion tests of endurance, WT males ran an average 62 min, whereas *Rdh10*+/− males ran an average 37 min (40% decrease), resulting in an equivalent 40% decrease in work accomplished (Figure 1B). WT females ran an average 33 min, i.e. ~47% less than WT males. In contrast *Rdh10*+/− females ran an average 72 min, ~2.2-fold longer running time than WT females and had the same % increase in work performed. Thus, loss of one *Rdh10* copy, and the accompanying moderate decrease in RA, reversed the relative running endurances of males and females.

### *Rdh10*+/− males and females differ in GM RA concentrations

To examine mechanisms, we focused on GM, because they perform predominantly in running, rather than standing, and have the most extensive overlap of pathways in mouse vs human (38). *Rdh10* mRNA in GM decreased sex specifically, declining by 53% and 33% in males vs females, respectively (Figure 1C). *Rdh10*+/− GM RA decreases of 38% and 17% in males and females, respectively, reflected the *Rdh10* mRNA decreases. Estrogen induces the retinol dehydrogenase *Dhrs9* (39–41). The mRNA of *Dhrs9* in male *Rdh10*+/− did not differ significantly from WT, whereas *Dhrs9* mRNA in female *Rdh10*+/− exceeded WT by 1.7-fold. Multiple other mRNAs of the retinoid metabolon, including the putative retinol dehydrogenase *Dhr7c*, the retinal dehydrogenases *Raldh1, 2, 3*, and the RA catabolic enzyme *Cyp26b1*, were unaffected in both sexes (Supporting Information Figure S3). The sex-specific differences in *Rdh10* and *Dhrs9* mRNAs would account for the difference between *Rdh10*+/− male and female RA concentrations and affect the physiological responses to Rdh10 reduction.

Because estrogen induces *Dhrs9*, we attempted to quantify estrogen concentrations in female sera. We were unable to detect estrogen in sera of female mice using an LC/MS/MS assay with a 2 fmol lower limit of detection (42). Estrogen concentrations in mice sera often occur below limits of detection, unless females are exposed to males. Instead, we quantified uterine weight, which reflects estrogen levels (43). *Rdh10*+/− uteri weighed 39% more than WT relative to total body weight, reflecting an estrogen increase (Figure 1D).

### *Rdh10*+/− males and females differ in myogenic transcription factor expression

mRNA levels of the myogenic transcription factor *Myog* (myogenin) increased 2.8-fold in *Rdh10*+/− male GM, relative to WT (Figure 2A). In contrast, *Rdh10*+/− female *Myog* mRNA decreased by 42%, relative to WT. mRNA of another myogenic transcription factor, *Myf5*, also increased in *Rdh10*+/− males (1.8-fold), whereas it decreased 30% in females. The mRNA of a third myogenic transcription factor *Myod* did not differ between WT and *Rdh10*+/− GM of either sex (Supporting Information Figure S4). To verify that mRNA levels represented protein levels, we performed western blots on Myf5 (Figure 2B,C). The data reflected adaptations in the mRNA. *Rdh10*+/− male Myf5 increased 3-fold, whereas *Rdh10*+/− female Myf5 decreased 38%. Other factors that regulate skeletal muscle mass and function include myostatin (Mstn), atrogin-1, and MuRF-1 (44–46). Mice deficient in Mstn have marked increases in skeletal muscle mass, because Mstn suppresses myogenesis by interfering with Myod. Striated muscle expresses atrogin-1 specifically. Atrogin-1 and MuRF are ubiquitin ligases that steer muscle protein to proteolysis. *Mstn* and *Murf1* mRNA did not change in *Rdh10*+/− of either sex relative to WT. The lack of differences in *Myod* in *Rdh10*+/− vs WT reflects the lack of differences in *Mstn* mRNA. *Atrogin-1* mRNA did not change in *Rdh10*+/− males relative to WT, but decreased 23% in *Rdh10*+/− females (Supporting Information Figure S5). Other genes associated with enhanced or diminished endurance in mice did not change including *Adcy5, Adrb2, Hif1a, Ppard, Pten, Scd1, Uchl1* (Supporting Information Figure S6) (47).

**Figure 2.**
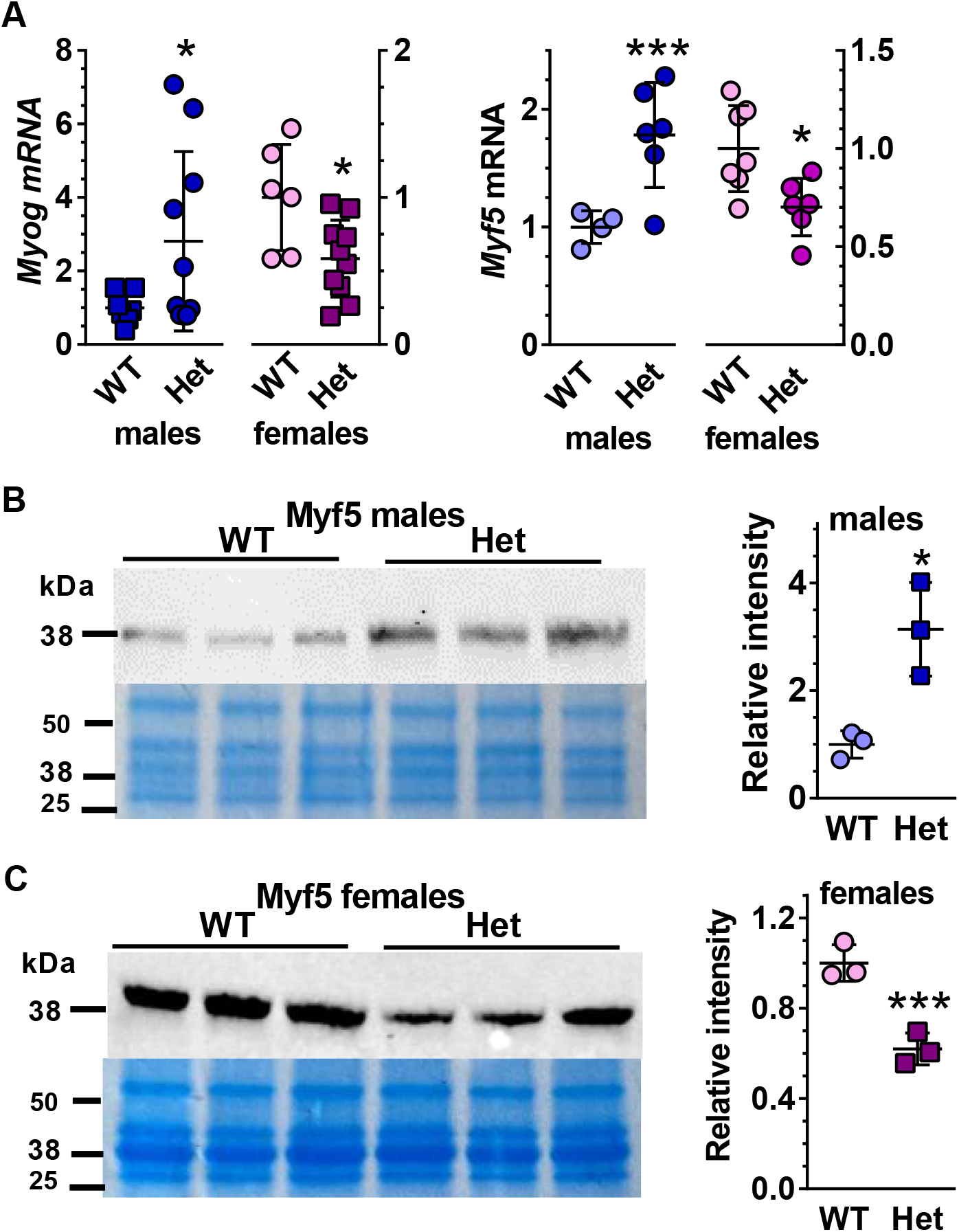
*Rdh10*+/− (Het) males and females differ from each other and WT in myogenic transcription factor expression. ***A***, mRNA of myogenic transcription factors *Myog* (n = 6 to 12) and *Myh5* (n = 4 to 12) in GM. ***B,C*,** Western blots of Myf5 in male and female GM, respectively. Data expressed as relative to WT (= 1) for each sex.

### Rdh10 affects GM fiber types in males

Differential myosin heavy chain (*Myh*) expression designates skeletal muscle fibers as type I (*Myh7*), type IIa (*Myh2*) or type IIb (*Myh4*) (48). Type I slow-twitch distance-running fibers contain abundant mitochondria. These fibers rely mainly on fatty acid oxidation. Fast-twitch “sprint” type IIb fibers have few mitochondria and rely mainly on glycolysis. Type IIa fibers rely on both oxidative and glycolytic pathways. GM consist of “mixed” fiber types, with ~54% type IIb fibers, and the rest distributed among types I and IIa/d. The mRNA of *Myh7*, an indicator of type I fibers, increased ~3-fold in *Rdh10*+/− male GM (Figure 3A). No changes occurred in *Rdh10*+/− *Myh2* (type IIa fibers) or *Myh4* (IIb fibers) mRNA in male GM. Immunostaining illustrates fibers types in *Rdh10*+/− male GM (Figure 3B). *Rdh10*+/− female GM *Myh7* mRNA remained the same as WT, as did *Myh2* and *Myh4* (Figure 3C). Immunostaining illustrates the nature of fibers in *Rdh10*+/− female GM (Figure 3D), Quantification of fiber numbers verified a 1.7-fold increase in *Rdh10*+/− male type I fibers and no change in female type I fibers (Figure 3E). Thus, *Rdh10*+/− males had decreased endurance despite compensation by increased slow-twitch oxidative fibers.

**Figure 3.**
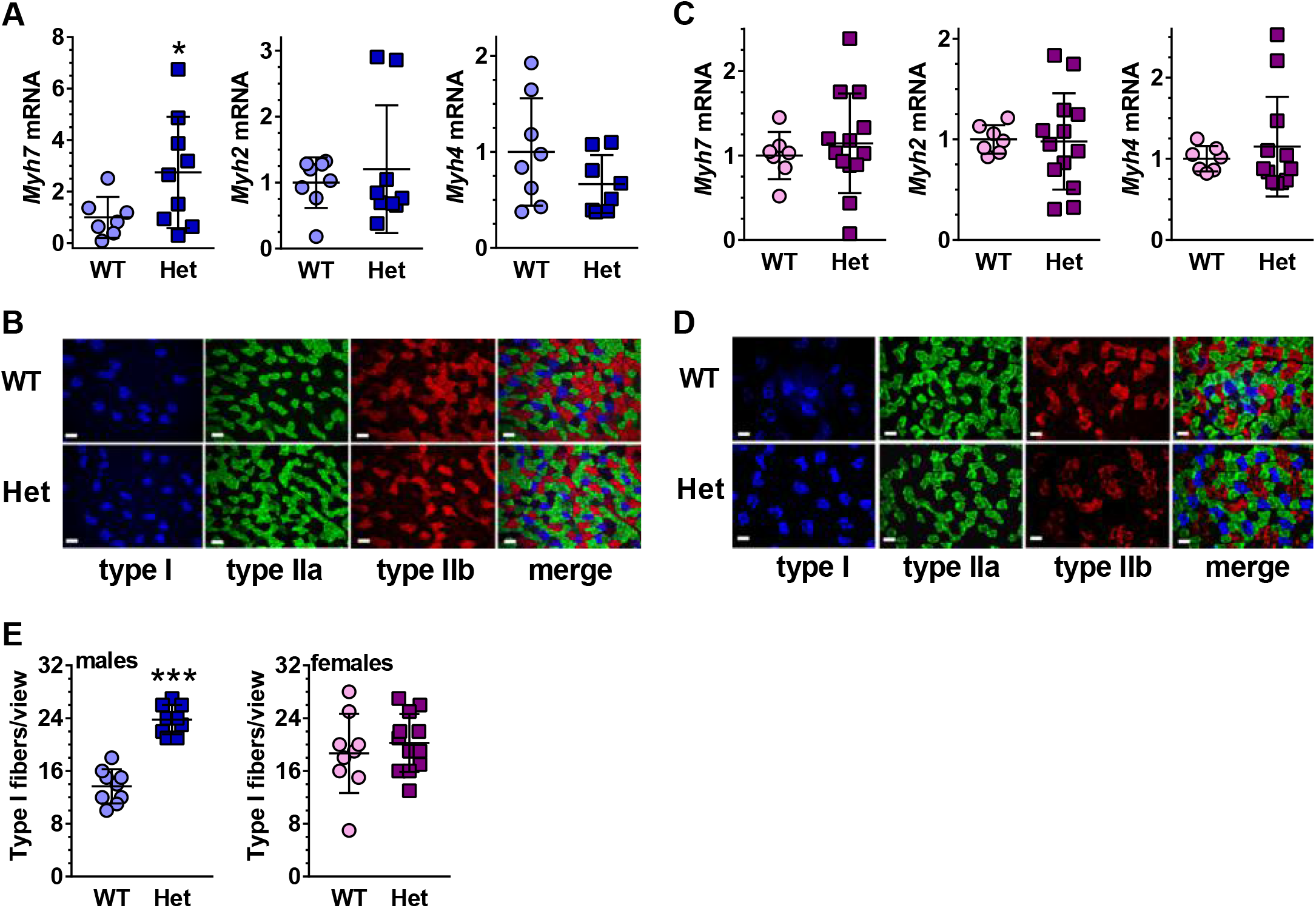
Rdh10 constrains the nature of GM fibers. *A*, mRNA of male *Myh7* (type I), *Myh2* (type IIa) and *Myh4* (IIb) genes (n = 7 to 9). *B*, Immunofluorescence of male GM fiber types. *C*, mRNA of female *Myh7, Myh2* and *Myh4* genes (n = 7 to 13). *D*, Immunofluorescence of female GM fiber types. *E*, Quantification of type I fibers (n = 9). Bars = 50 μm. Het, *Rdh10*+/−.

### *Rdh10*+/− males and females differ from each other and WT in muscle fiber characteristics

Normally, nuclei in muscle localize peripherally to fibers. Nuclei migration to the centers of fibers accompanies muscle dysfunction or regeneration (49). Figure 4A illustrates the difference between peripheral and centralized nuclei. Quantification showed that centralized nuclei increased 2.4-fold in *Rdh10*+/− male GM, but decreased 26% in *Rdh10*+/− female GM (Figure 4B). In addition, cross sectional areas (CSA) of *Rdh10*+/− GM fibers relative to WT adapted to a decrease in Rdh10 according to sex (Figure 4C). Numbers of larger male fibers increased

**Figure 4.**
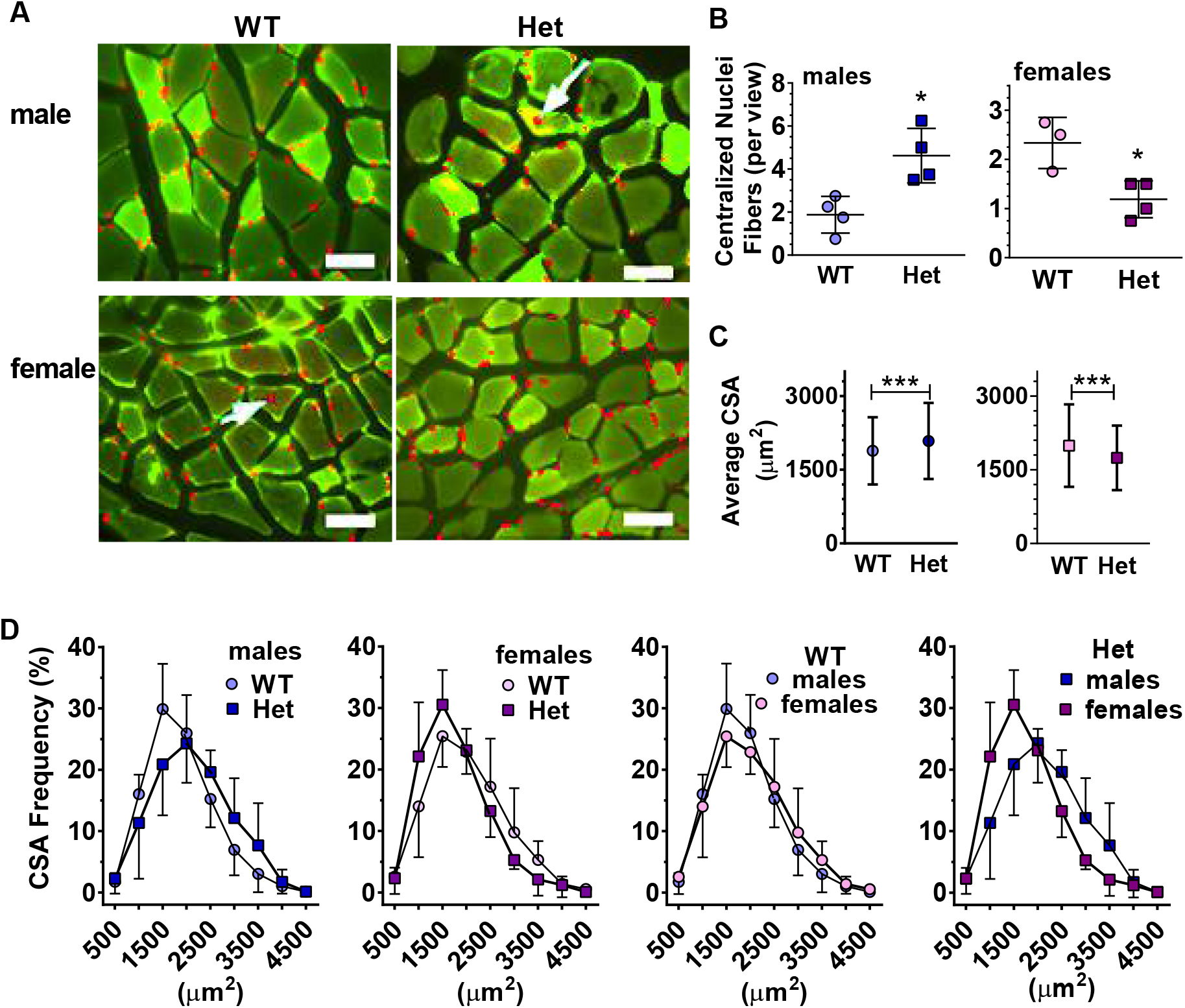
*Rdh10*+/− (Het) males and females differ from each other and WT in muscle fiber characteristics. *A*, Desmin immunostaining combined with DAPI staining of GM cryosections. The arrows denote centralized nuclei. Bars = 50 μm. Images were overexposed to allow visualization of nuclei. *B*, Quantification of average number of fibers with centralized nuclei per field of view using H&E staining (averages of 3 to 48 sections from each of 3 to 4 mice), *P<0.02. *C*, Average CSA/mouse (1454 to 1635 individual fibers quantified for each sex and genotype). *D*, The range of the CSA frequency distribution of data in *C*.

~10.7%, whereas numbers of larger female fibers decreased ~12.6%. The range of male CSA shifted to the right (larger) relative to WT (Figure 4D). The range of female CSA shifted to the left (smaller CSA). Direct comparison of WT male and female CSA revealed extensive similarity, indicating CSA normally do not diverge substantially according to sex (Figure 4E). All four curves (WT and *Rdh10*+/−, male and female) had similar if not identical areas, consistent with shifts in fiber size distributions in *Rdh10*+/−, rather than muscle growth or reduction. Direct comparison of *Rdh10*+/− male and female CSA illustrates the divergent impact of RA on CSA. Although *Rdh10*+/− female GM did not change in fiber type, nor show an increase in centralized nuclei, homogenization for the same length of time disrupted both *Rdh10*+/− male and female muscles much faster than WT (Supporting Information Figure S7). This indicates that the connective tissue surrounding the muscle fibers is weaker in *Rdh10*+/−, suggesting impaired tissue integrity in both sexes.

### *Rdh10*+/− males experience disturbances in GM lipid metabolism

Muscle weights relative to whole body weights also exhibited sexually dimorphic differences. *Rdh10*+/− male GM weighed 10% more than WT, whereas *Rdh10*+/− female GM weights did not differ significantly from WT (Figure 5A,B). The weight increase of *Rdh10*+/− males was not caused by an increase in protein content, as no marked differences occurred among GM protein amounts in either sex (Supporting Information Figure S8). Oil red O staining indicated greater TAG in *Rdh10*+/− males than other groups (Figure 5C). This was confirmed by quantification of TAG by biochemical assay, which revealed a 1.7-fold increase in *Rdh10*+/− male GM, but no increase in *Rdh10*+/− female GM relative to WT (Figure 5D). The decrease in Rdh10 triggered changes in mRNA of genes related to fatty acid biosynthesis, lipolysis, and mitochondria long-chain fatty acid import. Fatty acid synthase (*Fasn*), hormone sensitive lipase (*Hsl*), and carnitine palmitoyltransferase 1b (*Cpt1b*) mRNA decreased 30 to 40% in *Rdh10*+/− male GM (Figure 5E). *Rdh10*+/− female GM had a 50% increase in *Fasn* mRNA and a 36% decrease in *Cpt1b* mRNA, with no change in *Hsl* mRNA (Figure 5E). FASN protein reflected changes in mRNA, showing a 55% decrease in *Rdh10*+/− males and a 1.9-fold increase in *Rdh10*+/− females (Figure 5F,G,H). *Rdh10*+/− male and female GM mRNA did not change for *Atgl, CD36, Hk2, Mcad, Pgcla*, and *Ppara* (Supporting Information Figure S9). Insulin induces the elongation and initiation factor *Eif6*, which stimulates lipogenesis in liver and adipose, including transcription of *Fasn* (50). *Eif6*^+/−^ mice also endure reduced exercise endurance (38). Accordingly, consistent with insulin resistance, the decrease in FASN, and the decline in running endurance, *Eif6* mRNA decreased 26% in *Rdh10*+/− male GM, with no decrease in *Rdh10*+/− females (Figure 5I).

**Figure 5.**
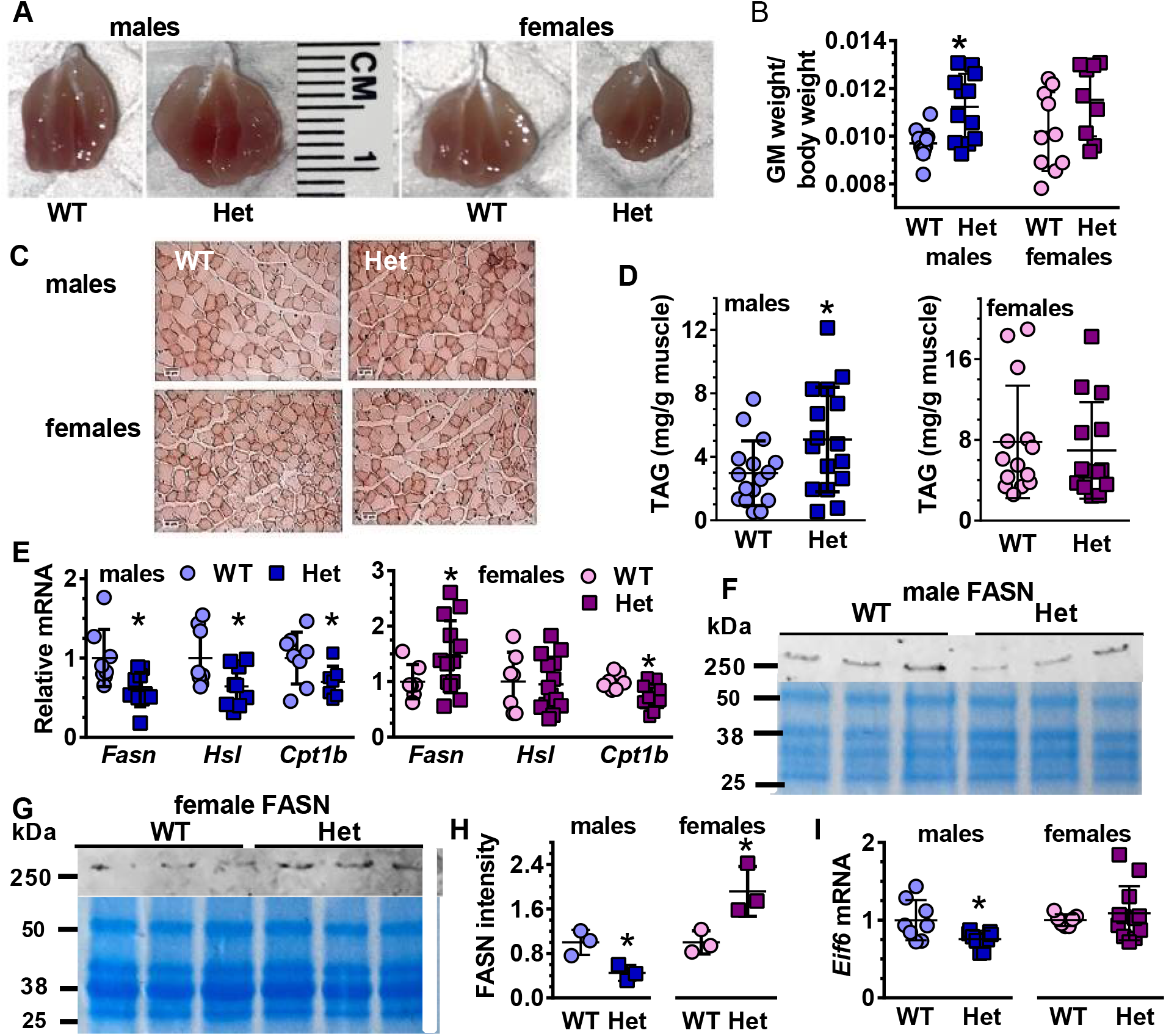
*Rdh10*+/− (Het) males experience disturbances in GM lipid metabolism. *A*, GM images. *B*, Quantification of GM weights relative to total body weights (n = 12). *C*, Images of oil red O staining. *D*, TAG content of GM (n = 14 to 16). *E*, mRNA of fatty acid metabolism related genes in male GM (n = 7 to 10) and female GM (n = 7 to 14). *F*, Western blots of FASN in male GM. *G*, Western blots of FASN in female GM. ***H***, Quantification of FASN western blots in GM. Data were normalized to the male WT signals set a 1. *I*, mRNA of *Eif6* in GM (n = 7 to 12).

### Glycogen and glucose adaptations differ sexually in *Rdh10*+/− GM

The glycogen content of *Rdh10*+/− GM did not differ from WT after fasting in either sex (Figure 6A,B). The mRNA of *Gys*, which encodes glycogen synthase, rate-limiting for glycogen synthesis, decreased 30% in *Rdh10*+/− male, but did not change in females (Figure 6C). The mRNA of *Pygm*, which encodes muscle glycogen phosphorylase, rate-limiting enzyme for glycogenolysis, did not change in *Rdh10*+/− of either sex. The lack of a decrease in muscle glycogen despite the decrease in *Gys*, likely reflects 16 h of fasting, which would deplete muscle glycogen.

**Figure 6.**
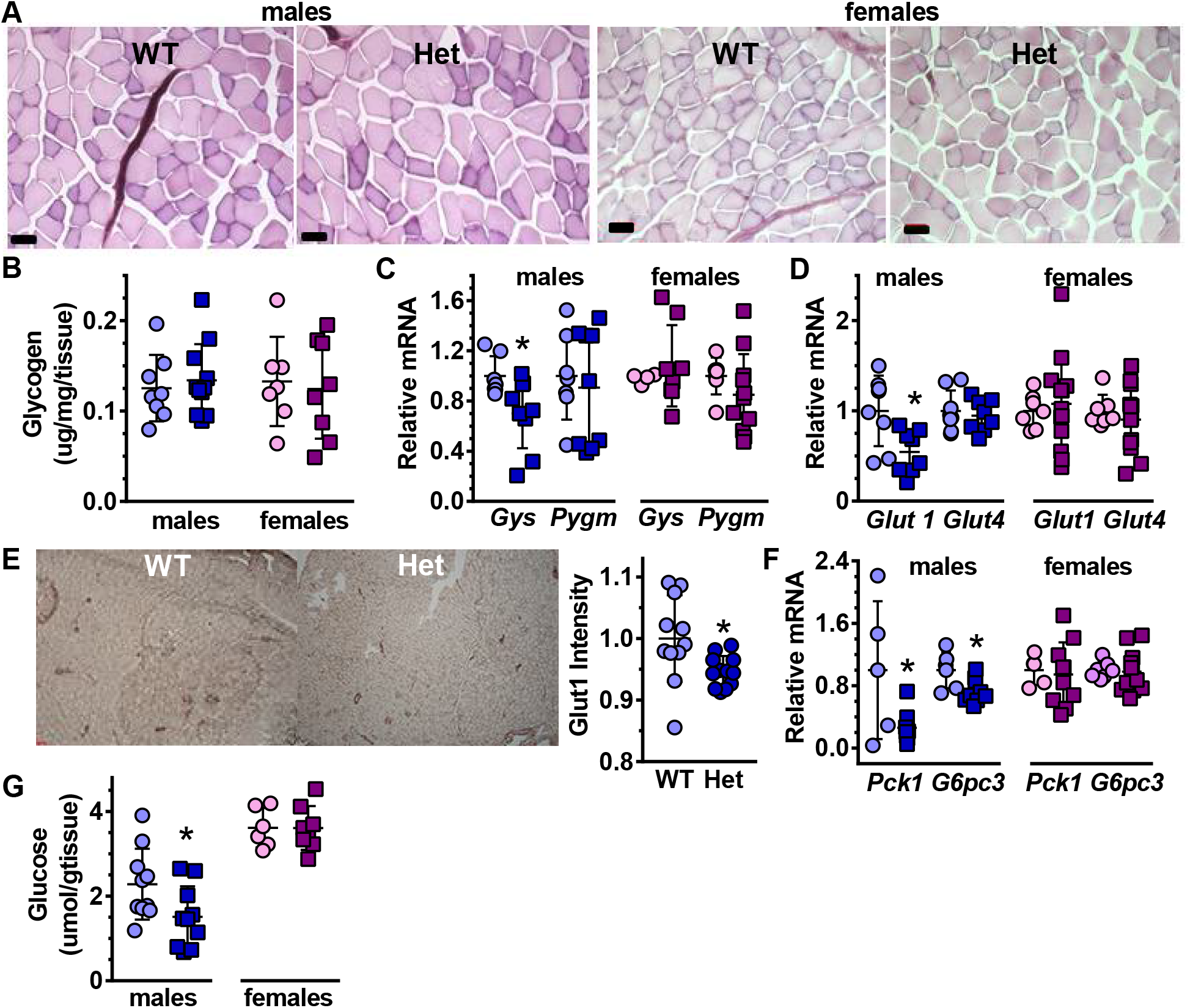
GM Glycogen and glucose adaptations by sex in *Rdh10*+/− (Het). *A*, Glycogen content (PAS staining, bars 50 μm). *B*, Quantification of *A* (n = 7 to 11). *C, Gys* and *Pygm* mRNA (n = 7 to 11). *D*, GM *Glut1* and *Glut4* mRNA (n = 7 to 13). *E*, Glut1 immunostaining in males (bars 200 μm). Quantification was done on 4 views for each of 3 mice/genotype. *F, Pck1* and *G6pc3* mRNA (n = 5 to 10). *G*, Glucose content (n = 6 to 8).

Muscle relies for glucose uptake on the glucose transporters Glut1 and Glut4. *Glut1* mRNA diminished 36% in *Rdh10*+/− male GM, but did not change in *Rdh10*+/− females (Figure 6D). *Glut4* mRNA did not change in *Rdh10*+/− of either sex. The intensity of Glut1 immunostaining decreased a modest 5%, which could have a profound influence on glucose uptake because the mice were fasted 16 h (Figure 6E). The mRNA of two key genes that encode enzymes essential to gluconeogenesis, *Pck1* (phosphoenolpyruvate carboxykinase 1) and *G6pc3* (glucose-6-phosphatase catalytic subunit 3) decreased by 73 and 27%, respectively, in male GM, but not in female GM (Figure 6F). The average amount of glucose in fasted male GM decreased by 34%, reflecting the decrease in *Glut1* and gluconeogenesis genes, and the glucose intolerance and insulin resistance of male Rdh10+/− mice (Figure 6G).

### *Rdh10*+/− male GM suffer reduced mitochondria function; females gain function

We assessed mRNA of genes associated with GM mitochondria function to determine the extent of electron transport chain impairment. mRNAs of complex IV components (cytochrome C oxidase) *Cox5a* and *Cox8b* declined by 46% and 39%, respectively, in *Rdh10*+/− male GM (Figure 7A). *Rdh10*+/− female GM *Cox5a* and *Cox8b* mRNA increased from 20 to 36%, but data were shy of statistical significance at P < 0.05. The mRNA amounts of other electron transfer chain components in *Rdh10*+/− GM did not differ from WT (Supporting Information Figure S10). These included *Ndufs2* (complex I), *Sdhb* (complex II), *Uqcrc2* (complex III), *Cycs* (cytochrome c), and *Apt5al* (complex V). Cox5a protein expression in GM lysates decreased in *Rdh10*+/− male GM by an average 28%, followed by a complex IV activity decrease of 13% and a fasting ATP decrease of 60% (Figure 7B,C). In contrast, Cox5a relative protein expression increased by an average 40% in *Rdh10*+/− female GM, accompanied by a 40% increase in complex IV activity, and a >3-fold increase in fasting ATP (Figure 7B,D).

**Figure 7.**
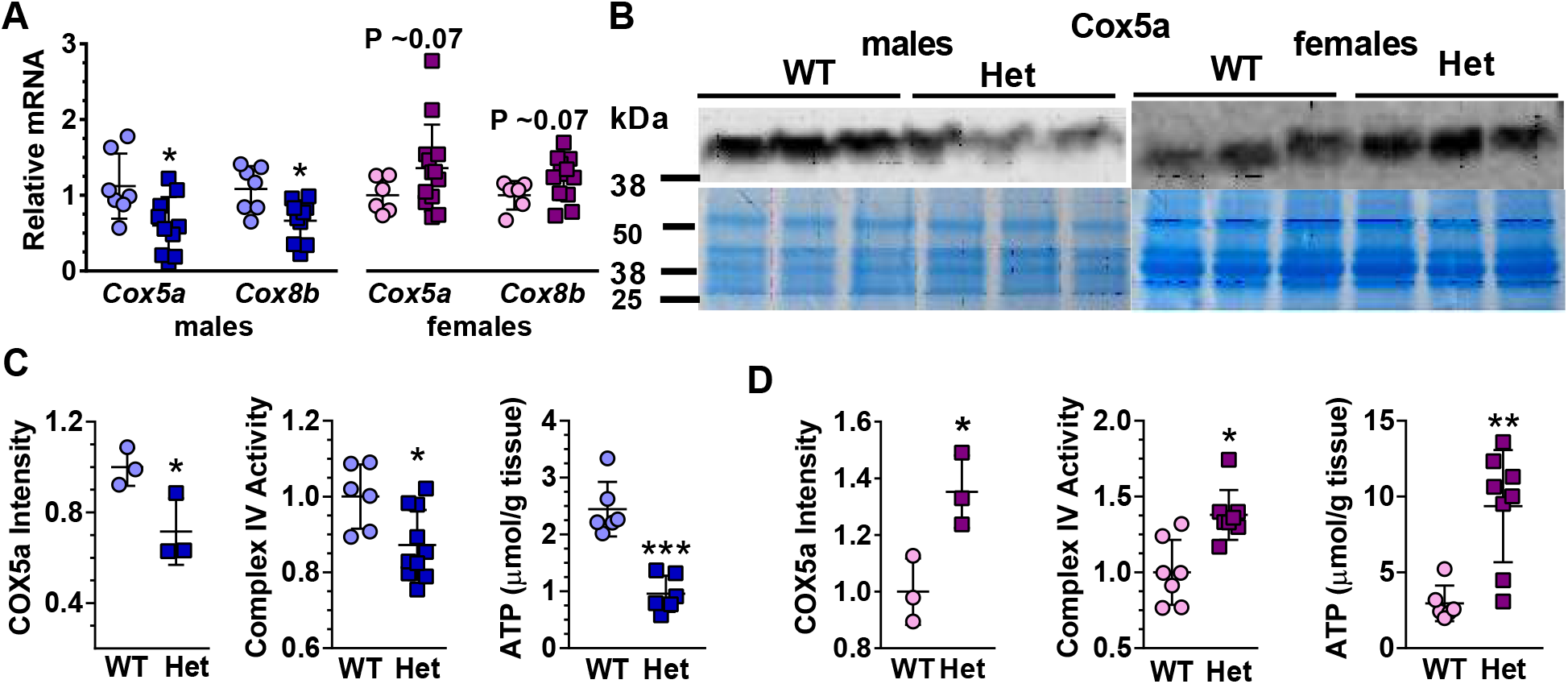
*Rdh10*+/− (Het) male GM suffer reduced mitochondria function; females gain function. *A*, Electron transport chain complex constituents *Cox5a* and *Cox8b* mRNA in GM (n = 6 to 13). *B*, Western blots of Cox5A in GM. ***C*,** Quantification of male GM data in *B* relative to WT, relative complex IV activity (n = 6 to 10), and ATP concentration (n = 6 each genotype). *D*, Quantification of female GM data in B relative to WT, relative complex 4 activity (n = 7 to 8), and ATP concentration (n = 6 to 8).

## Discussion

We reported that eliminating one *Rdh10* copy in vivo (*Rdh10*+/−) reduced RA modestly in adult liver and white adipose (≤ 25%), and increased diet-induced obesity in male mice (20). Only males endured liver steatosis when fed a high-fat diet, and only females sustained increased bone marrow adipocyte formation, regardless of dietary fat. Here we report the impact of marginally reduced RA in muscle, showing an outcome influenced by increased estrogen in females, associated with an attenuated decrease in RA relative to males. Assessing the whole body *Rdh10* heterozygote enabled exposure of estrogen-driven differences in RA function, and permitted assessment of overall RA impact on adiposity, insulin resistance and carbohydrate vs. fat use as fuels.

Male C57/Bl6 mice have larger body weights and more muscle mass compared to age-matched females (20). This difference in lean body mass, along with a greater reliance on fatty acid oxidation (RER), enables longer running times in WT males. The modest but significant decrease in *Rdh10*+/− male reliance on fatty acid oxidation and increased reliance on carbohydrates, also indicated by the RER, would affect running endurance, reflecting decreases in GM *Cpt1b* and *Hsl* mRNA, driving the increase in GM TAG. The decrease in *Gys* and *Glut1* would reduce glucose uptake and storage in male GM, contributing to whole-body glucose intolerance, as muscle accounts for >70% of insulin-stimulated glucose use (51). Decreases in *Pck1* and *G6pc3* would contribute to the decrease in male GM glucose. These decreases in energy availability, reflected in glucose intolerance and insulin resistance, plus impairment of the electron transport chain, as indicated by decreases in RA-induced cytochrome C oxidase subunits, would hinder ATP production in male GM and deny energy for running endurance (52). Impaired energy use would prompt insulin resistance, which explains the decrease in *Fasn*, an insulin-regulated gene. The increase in type I (oxidative) fibers suggests a futile attempt at compensation for diminished fatty acid oxidation and running endurance.

The increased dependence on fatty acid oxidation in *Rdh10*+/− females (RER) is consistent with increased endurance, as is a lack of glucose intolerance or insulin resistance, indicated by GTT and ITT, as well as increased *Fasn* mRNA. The increased reliance on fatty acid oxidation, and the increase in cytochrome C oxidase subunits, along with more efficient glucose uptake than males, produced more ATP and greater running endurance. These data show that RA has a complex impact on post-natal skeletal muscle function that alters ATP generation via both oxidative and non-oxidative mechanisms. These sexually dimorphic differences in muscle contribute to the whole-body phenotype of *Rdh10*+/−.

The surprising increase in female muscle performance with a decrease in RA would seem to challenge concepts of RA function, because decreased RA usually diminishes cell, organ and physiological performance (53). Then again, this outcome strengthens insight into interactions between RA function and estrogen concentrations, and indicate that estrogen enhances RA signaling, in addition to inducing its biosynthesis. Enhancement of RA action by estrogen and impairment of estrogen action by RA has been noted in cell lines. Estrogen sensitizes human breast cancer cells to RA by inducing RARβ (54). Conversely, RA restricts estrogen receptor α action in the estrogen-sensitive human breast cancer cell line MCF7/BUS (55), and inhibits estrogen-induced proliferation of MCF-7 cells (56). RA also promotes homeostatic maintenance of the adult mouse uterus, preventing excess proliferation (57). Notably, estrogen action in muscle increases insulin sensitivity, prevents lipid accumulation and promotes metabolic health (58). These data indicate that estrogen contributes to improving the impact of reduced Rdh10 in females through effects on the RA concentrations and function. In contrast to estrogen actions, androgens repress RAR mRNA in rat prostate (59), whereas RA suppresses androgen receptor expression in rat testis (60). The decreased performance by *Rdh10*+/− males would reflect decreased RA signaling and amplified repression of RAR by androgen.

A multifaceted process controls skeletal muscle development, regeneration and function (61). The myogenic transcription *Myog* regulates transcription in muscle. Ablating *Myog* produces a mouse with enhanced performance in treadmill running, as a result of improved fuel metabolism (62). The decreased running performance of *Rdh10*+/− males and the increased performance of *Rdh10*+/− females, and their decrease and increase in fuel metabolism, reflects the increase and decrease, respectively, of *Myog*. The myogenic factors, Myf5 and MyoD commit stem cells to a skeletal muscle lineage. RA effects on Myf5 and MyoD have been studied during embryogenesis, but post-natal effects of endogenous RA have not been investigated (63). Cumulative data in cell lines and chick limb buds indicate concentrationdependent RA effects on myogenesis (64–66). These results may reflect retinoid-related hormesis: i.e., RA effects depend on concentrations (67). Beneficial effects occur as concentrations increase to an optimum, which presumably occurs at the low nM RA that occurs in tissues (68). As concentrations increase beyond an optimum, beneficial effects subside and pharmacological effects ensue; ultimately toxic effects prevail.

Hormesis also may contribute to sexual dimorphism. Estrogens vs. androgens exert different effects on RA functions apparently by increasing or decreasing RAR expression, respectively, which would impact the RA dose-response curves based on sex. The sex-hormone/RA nexus would help explain how modest alterations in RA have contrary sex-related effects. Moreover, RA effects on myogenic differentiation depend on the exact cellular context, as location-specific interactions with the local chromatin environment affects transcriptional activity (69), as well as expression of distinctive retinoid binding-proteins (70).

## Conclusions

The present data complement insight into the anti-obesity effects of RA by revealing that endogenous RA promotes skeletal muscle fatty acid oxidation, glucose uptake, mitochondrial function and ATP biosynthesis in males, but restricts the same in females. These male and female specific actions also affect glucose tolerance and insulin resistance. Contrasting interactions between estrogens vs androgens on RA biosynthesis and function suggest mechanisms for these dimorphic effects. This sexually dimorphic insight into RA in muscle contributes to understanding differences between males, pre-menopausal females, and postmenopausal females with respect to regulation of metabolic health. Given the complexity of RA actions in vivo and its hormesis-dependent effects, partial ablation of *Rdh10* should continue to provide opportunity to distinguish physiological from pharmacological and/or toxic actions of RA.

## Experimental procedures

### Mice and diets

Mouse experimental protocols were approved by the University of California-Berkeley Animal Care and Use Committee. *Rdh10*-floxed mice were bred with mice expressing CMV-Cre (B6.C-Tg (CMV-Cre) 1Cgn/J) to generate a whole-body knockout of *Rdh10* as described (20). Homozygous *Rdh10* knockout causes embryonic lethality, therefore *Rdh10* heterozygotes (*Rdh10*+/−) were studied. CMV-Cre+ littermates served as controls (WT). Mice were backcrossed into the C57BL/6J background >12 generations. Mice were fed an AIN93G diet containing 4 IU/g vitamin A with 50% fat since weaning and were fasted overnight prior to tissue collection to promote fatty acid use, unless otherwise noted. Mice were 4 to 5-months old.

### Respiratory exchange ratio (RER)

RERs were determined with a Columbus Instruments Comprehensive Laboratory Animal Monitoring System (CLAMS). Mice were acclimatized to chamber cages 24 h before values were recorded. The CLAMS was housed in a temperature controlled (23 °C), 12-h light/dark cycle room, with lights on at 7 AM. Mice had access ad libitum to food and water.

### Biochemical assays

ATP was quantified with an ATP Colorimetric/Fluorometric Assay Kit (Biovision K354). TAG concentrations were determined using a Triglyceride Colorimetric Assay Kit (Cayman 10010303). Glycogen contents were measured with a Glycogen Colorimetric/Fluorometric Assay Kit (Bio Vision K646). Complex IV activity was analyzed with a Complex IV Rodent Enzyme Activity Microplate Assay Kit (Abcam ab109911). Glucose was quantified with a Promega Kit (Glucose-Glo^™^ Assay).

### Run-to-exhaustion test

We used a motor-driven treadmill (Columbus Instruments, Exer-6M Open Treadmill) to assess running endurance. To encourage mice to run, tactile stimuli were provided by a manual light tap with a small paint brush to the hindquarters when the animal reached the back of the treadmill. On days 0 and 1, mice were placed on the treadmill to acclimate for 10 min with a speed of 10 m/min at 0 % gradient. On day 2, running capacity was determined by placing mice on the treadmill without treadmill motion for 2 min for acclimation. Next, mice were subjected to a 5 min warm-up period at 10 m/min. Speed was then increased by 2 m/min each 2 min until a maximum of 20 m/min. Mice were then run to exhaustion, defined as sitting at the back wall for 10 s despite continual tapping with the brush. Work (Joules) was calculated from body weight (kg) x 9.8 x distance run in m.

### Antibodies

Primary antibodies: Myh2 (3 μg/ml, Developmental Studies Hybridoma Bank, sc-71, AB_2147165), Myh4 (3 μg/ml, Developmental Studies Hybridoma Bank, BF-F3, AB_2266724), Myh7 (3 μg/ml, Developmental Studies Hybridoma Bank, BA-F8, AB_10572253), Desmin (1:20. Thermo Fisher Scientific, MA5-13259), Cox5a (1:1000, Abcam, ab110262, AB_10861723), Glut1 (1:200, Abcam, ab652, AB_305540), Myf5 (1:10000, Abcam, ab125078, AB_10975611), Fasn (1:1000, Abcam, ab22759, AB_732316). Secondary antibodies: goat-anti-mouse IgG2b Alexa 350 (1:200, Thermo Fisher A-21140, AB_2535777), goat-anti-mouse IgG1 Alexa 488 (1:200, Thermo Fisher A-21121, AB_2535764), goat-anti-mouse IgM Alexa 555 (1:200, Thermo Fisher A-21426, AB_2535847), IRDye^®^ 800CW goat anti-mouse IgG Secondary Antibody (1:10000, Licor, 926-32210, AB_621842), IRDye^®^ 800CW goat antirabbit IgG Secondary Antibody (1:10000, Licor, 926-32211, AB_621843), Anti-rabbit IgG HRP-linked Antibody (1:200, Cell Signaling Technology, 7074P, AB_2099233).

### Histology and Immunofluorescence

Entire GM were dissected from legs, snap-frozen in liquid nitrogen and stored at −80 °C. Cryosections (10 μm each) were harvested from frozen muscles with a Cryostat (Leica CM3050) and processed for histological and immunofluorescence analysis. Sections were stained with hematoxylin (Sigma GHS3) and eosin Y (RICCA 2845-32), washed with distilled water, and mounted in Shur Mount (General Data LC-W). For cross sectional area (CSA) quantification, all fibers in each view were quantified from 4 mice per group with 4 sections for each mouse, except for one of the female WT and one female Het for which 3 sections were analyzed. In total 1454 to 1635 fiber sections were quantified for CSA. Oil red O was used to stain lipids (34). Sections were mounted in Shur Mount and glycogen was visualized by Periodic Acid-Schiff staining (PAS) (MilliporeSigma 395B-1KT) following manufacturer’s instructions. Slides were imaged with a Zeiss AxioObserverZ1 microscope at 40x magnification, using QImaging 5MPx a MicroPublisher color camera (Biological Imaging Facility, UC-Berkeley).

For immunofluorescence imaging, sections were warmed to room temperature for 40 min and washed three times with 0.1% PBST. After blocking with 5% NGS/PBST (normal goat serum in 1% Tween 20/phosphate-buffer) 1 h at room temperature, sections were incubated in 5% NGS/PBST with a mixture of three primary antibodies against Myh7 (IgG2b), Myh2 (IgG1) and Myh4 (IgM), obtained from DSHB at the University of Iowa. After three 5 min washes in PBST, sections were incubated 1 h at room temperature with a mixture of three goat anti-mouse secondary antibodies against IgG2b (Alexa 350), IgG1 (Alexa 488), and IgM (Alexa 555) in 5% NGS/PBST, followed by three washes in PBST. For analysis of centralized nuclei, sections were first incubated in the primary anti-Desmin monoclonal antibody overnight at 4 °C. For nuclear staining, sections were incubated in 300 nM DAPI (4’6-diamidino-2-phenylindole, Thermo Fisher Scientific, D1306) working solution for 5 min at room temperature, followed by washing in PBS. Sections were mounted with Prolong Diamond Anti-fade mountant (Thermo Fisher P36965) and sealed with colorless nail polish. Washing and blocking for IHC staining were the same as for immunofluorescence. Slides were incubated overnight at 4°C with primary glucose transporter 1 antibodies (Glut1), washed with 0.1% PBST, and incubated with goat-anti-rabbit IgG HRP. Stained slides were examined at 40 x magnification with oil immersion using Zeiss Axiovert M1 fluorescent microscope equipped with Hamamatsu Orca CCD camera and CoolLED pE-300 white LED illuminator, and Zeiss Z1 AxioObserver inverted microscope equipped with a 40 x immersion lens, X-Cite 120 LED fluorescence source and Qimaging Retiga SRV CCD camera (Biological Imaging Facility, UC Berkeley). To quantify Glut1 intensity, the reciprocal of brightness in the blue channel-measure was used in Image J.

### RNA extraction and qPCR

Total RNA was extracted from tissues with TRIzol reagent (Life 15596026). The RNA concentration was measured with a Nanodrop spectrophotometer and reverse-transcribed with iScript (Bio-Rad 1708841), followed by qPCR reactions using Prime Time Gene Expression Master Mix (Integrated DNA Technologies 1055771). qPCR data were normalized to the housekeeping genes *Gusb* and *Tbp*. Each *Rdh10*+/− mRNA was normalized to its WT littermates. We used pre-designed primers purchased from Integrated DNA technology. Primer details are provided in Supporting Information (Table 1).

### Western blots

Immunoblotting was performed as described (35). Muscles were thawed on ice and homogenized with a TissueLyser II (Qiagen Inc.) in 1 mL cold RIPA buffer (Thermo Fisher 89900) supplemented with protease inhibitor (Thermo Fisher A32965) and phosphatase inhibitor (Roche 4906845001). Tissue lysates were denatured and centrifuged. Protein was quantified in the supernatant with the BCA protein assay (Thermo Fisher 23225), subjected to SDS-PAGE and transferred onto PVDF membranes (Bio-Rad 1620177). Membranes were blocked with 5% fat-free milk, incubated with primary antibodies overnight (4 °C), and with secondary antibodies 1 h at room temperature. Membranes were scanned and signal intensities were quantified using a LiCOR Odyssey imager. Protein loaded per lane was determined with Coomassie blue staining (Thermo Fisher 46-5034) and analyzed with NIH Image J. Data were normalized to total protein loaded per lane. Units are arbitrary (AU), normalized to WT males set to 1, unless noted otherwise.

### RA quantification

RA was quantified by LC/MS/MS as published, except methanol was used instead of ethanol, and homogenates were centrifuged at 1,200 x g for 5 min to remove particulates (36). LC was done as published (37).

### Statistics

All data are present in this manuscript. Data were plotted as means ± SD, unless noted otherwise. Two-tailed unpaired t-tests were used to analyze results between two groups. Welch’s corrections were applied to groups without equivalent SD. Significantly different from WT: *P < 0.05, ** P < 0.01, ***P < 0.001, unless noted otherwise; n = number of mice per genotype/sex.

## Acknowledgements

This work was supported in part by NIH grants DK102014 (JLN) and DK112754 (JLN), and Sponsored Projects for Undergraduate Research supported by the College of Natural Resources, UC-Berkeley (JLN). The content is solely the responsibility of the authors and does not necessarily represent the official views of the National Institutes of Health.

Authors are grateful to Milena Tintcheva for generating preliminary data during early investigations, and for maintaining the mouse colony, and to Delphine Gaubert for help with some experiments. Imaging was conducted at the College for Natural Resources Biological Imaging Facility. Authors are grateful to Dr. Steve Ruzin and Dr. Denise Schichnes for exceptional support of this project with imaging expertise and help with imaging data collection.

## Author contributions

Y. Zhao, M. Vuckovic and J.L. Napoli designed the research. Y. Zhao, M. Vuckovic, H.S. Yoo, N. Fox, A. Rodriguez, and K. McKessy conducted experiments. Y. Zhao, M. Vuckovic, H.S. Yoo, N. Fox, A. Rodriguez, K. McKessy and J.L. Napoli analyzed data. Y. Zhao and J.L. Napoli wrote the initial draft of the manuscript. M. Vuckovic and Y.S. Yoo edited the manuscript.

## Conflict of interests

The authors declare that they have no conflicts of interest with the contents of this article.

## Abbreviations

CLAMS: Comprehensive laboratory animal monitoring system
CSA: cross sectional areas
GM: gastrocnemius muscle
RA: all-*trans*-retinoic acid
RER: respiratory exchange ratio
*Rdh10*+/−: *Rdh10* heterozygote
TG: triacylglycerol
WT: wild type

## Supporting Information

**Figure S1.**
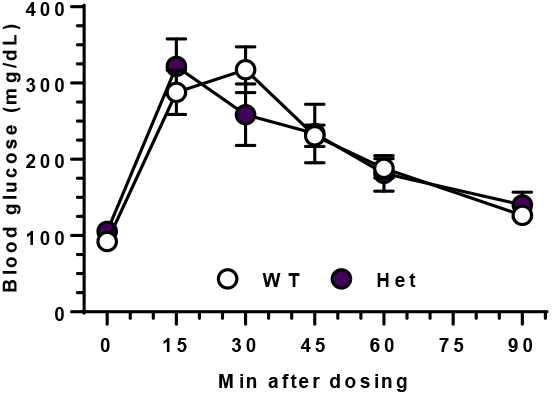
GTT of females. Mice were fasted 16 h and then dosed i.p. with 1.5 g glucose/kg body weight. Six to 8 mice/genotype. Blood glucose from tail tips was measured with a glucometer.

**Figure S2.**
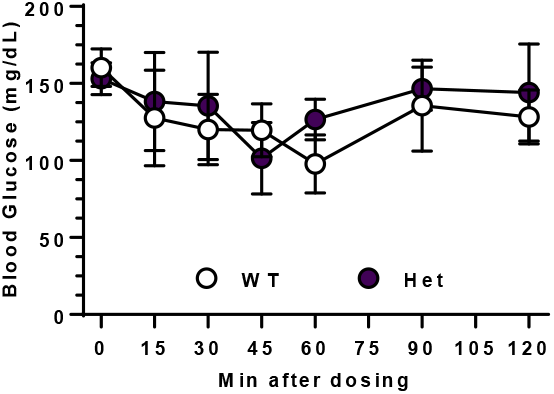
ITT of females. Mice were fasted 4 h and placed in individual cages without food, but with free access to water. Insulin (0.5 IU/kg body weight) was injected i.p. into 4 to 5 mice/genotype. Blood glucose from tail tips was measured with a glucometer.

**Figure S3.**
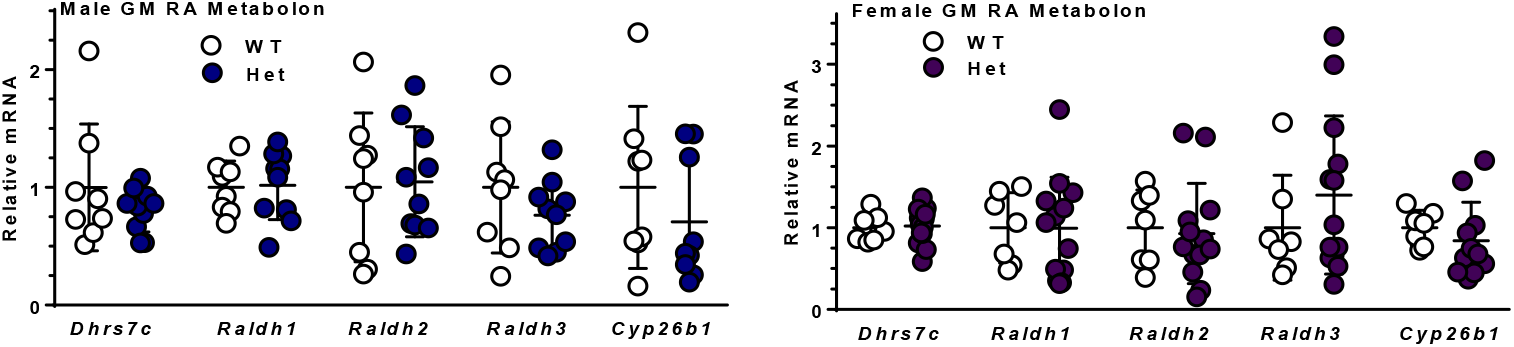
mRNA of the retinoid metabolon genes not changed in Het. Seven to 13 mice per gene/genotype/sex. Muscle does not express *Rdh1* mRNA.

**Figure S4.**
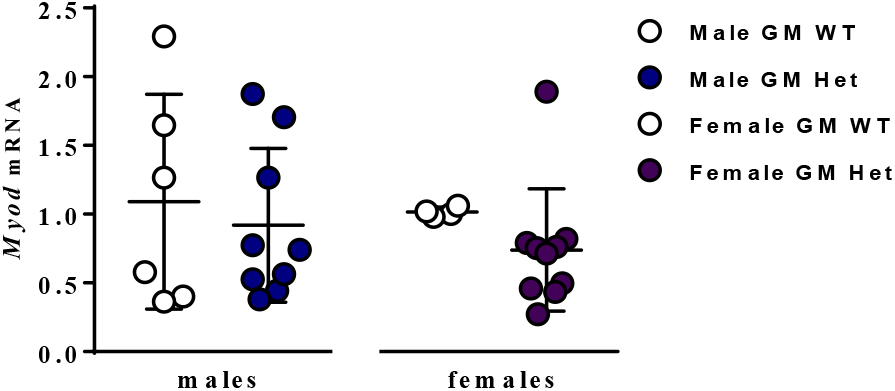
qPCR of *Myod* mRNA. Four to 10 mice per genotype/sex.

**Figure S5.**
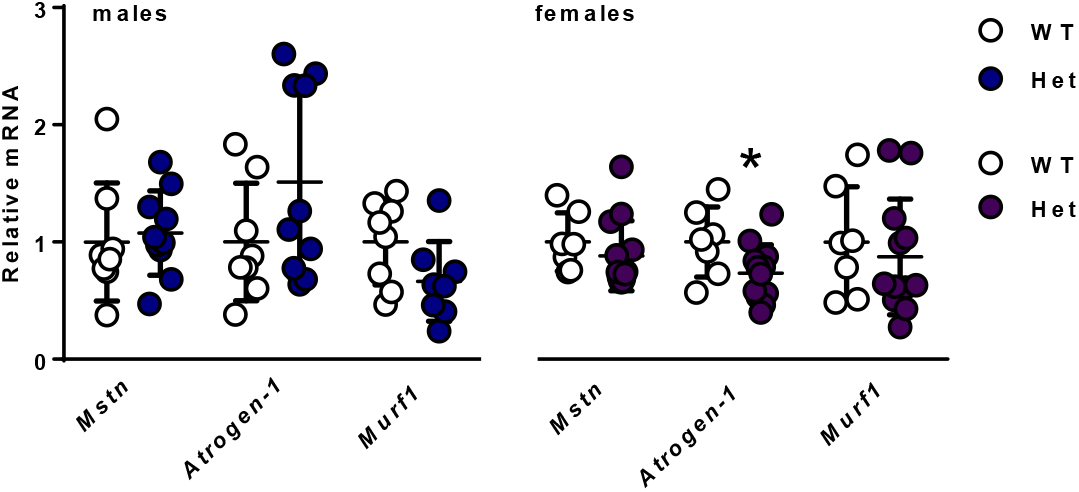
mRNA of genes associated with skeletal muscle mass and function. Seven to 13 mice gene/geneotype/sex.

**Figure S6.**
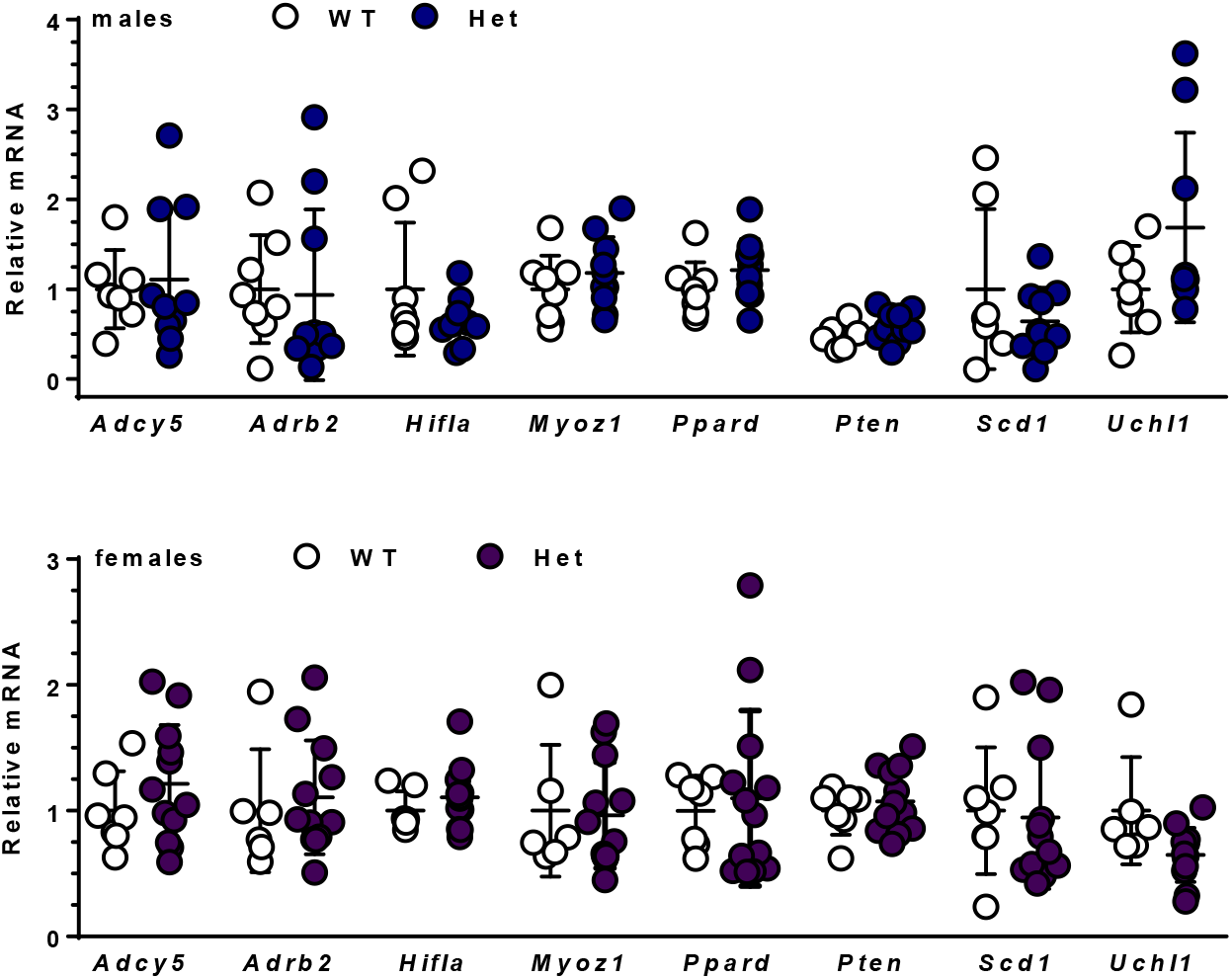
mRNA of genes that affect running endurance in mice. Four to 10 mice per gene/genotype/sex.

**Figure S7.**
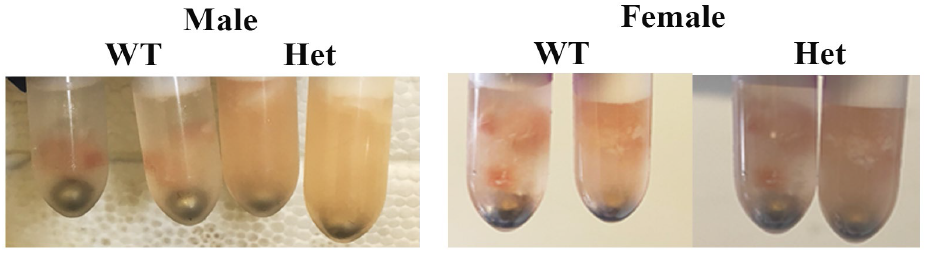
Representative images of GM muscle homogenization. Entire muscles were homogenized with Tissuelyser II (Qiagen) at 30 Hz for 15 min. WT of both sexes remained largely intact, whereas Het of both sexes were substantially disrupted under the same conditions.

**Figure S8.**
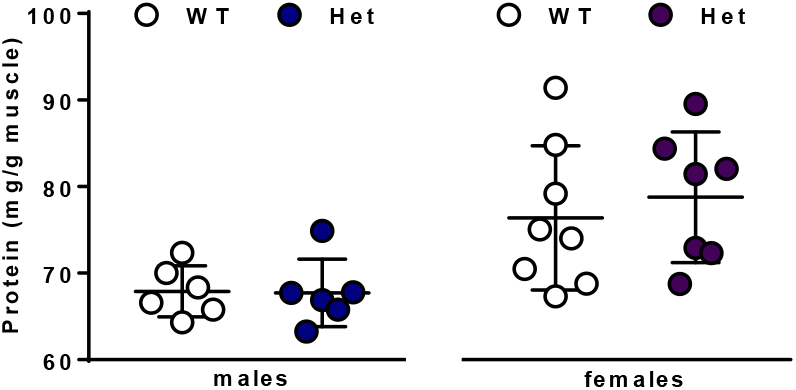
Total protein amounts in GM. Six to 8 mice per muscle/genotype/sex.

**Figure S9.**
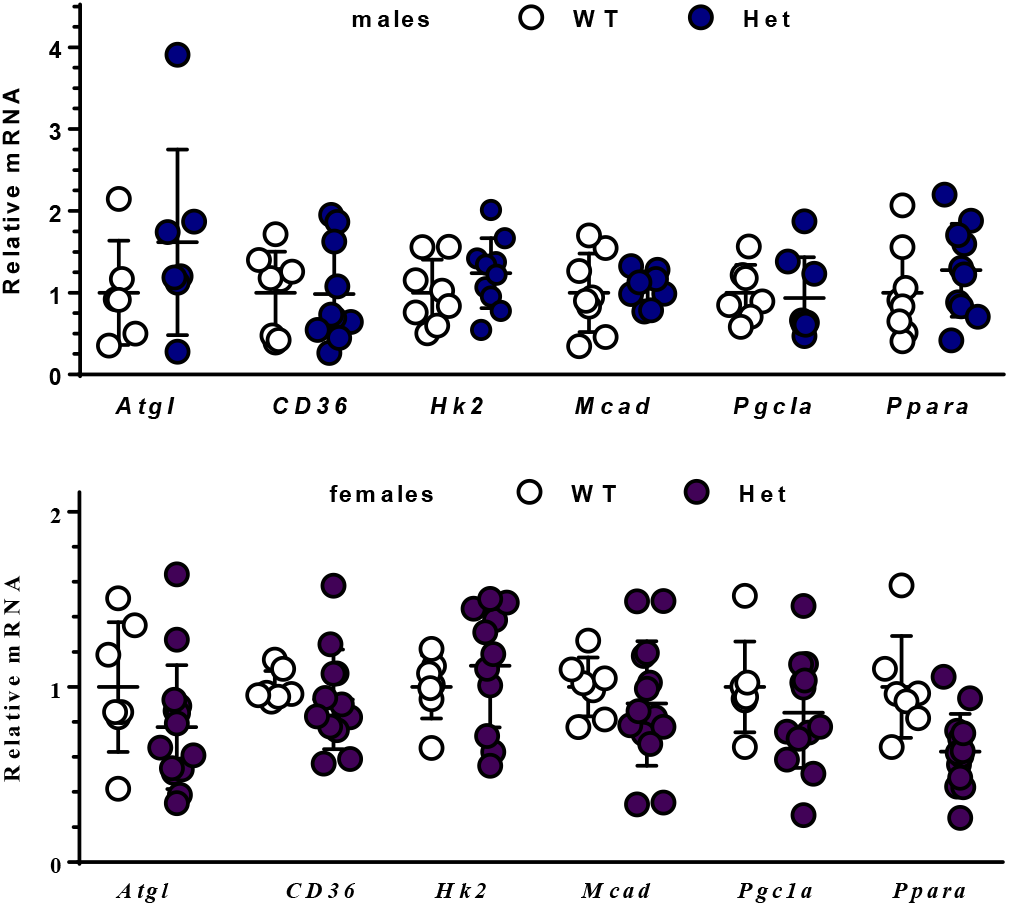
mRNA of fatty acid metabolism genes in GM. Six to 14 mice per gene/genotype/sex

**Figure S10.**
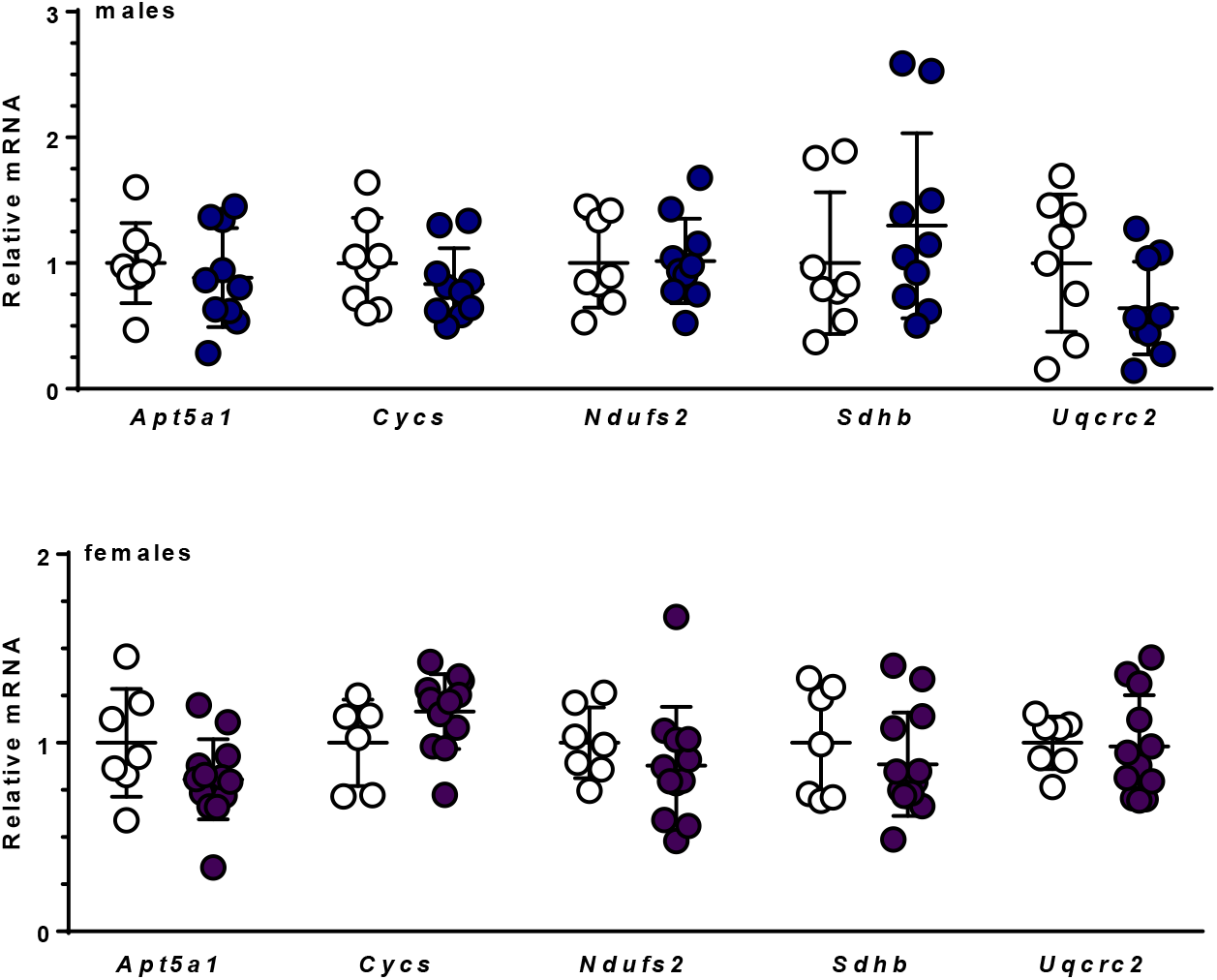
Mitochondria function genes. Six to 12 mice per gene/genotype/sex/muscle.

**Supporting Table 1.**
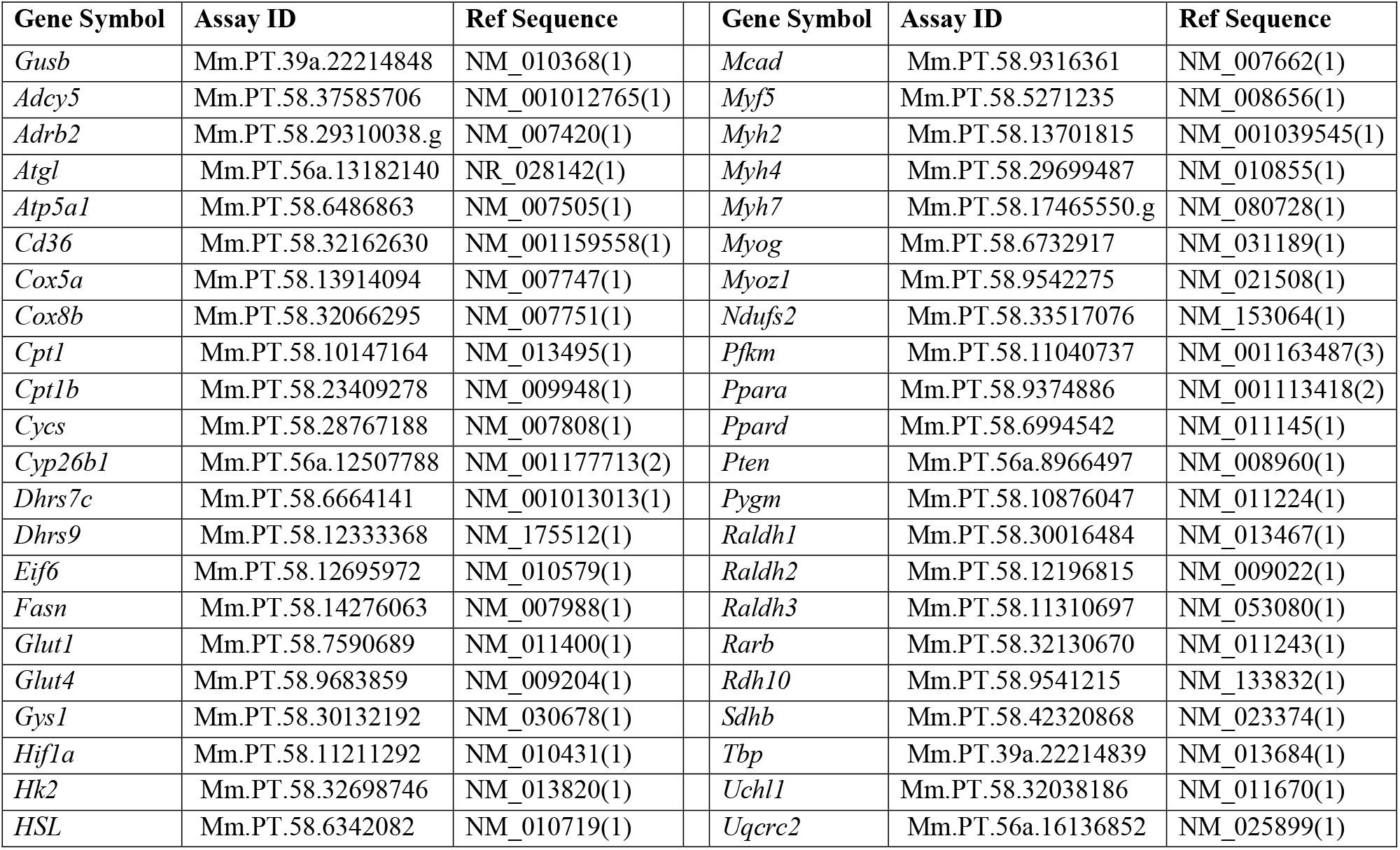
Primers used for qPCR.

## REFERENCES

1. Iskakova, M., Karbyshev, M., Piskunov, A., and Rochette-Egly, C. (2015) Nuclear and extranuclear effects of vitamin A. Can. J. Physiol. Pharmacol. 93, 1065–1075

2. Rhinn, M., and Dollé, P. (2012) Retinoic acid signaling during development. Dev. Camb. Engl. 139, 843–858

3. Nagpal, I., and Wei, L.-N. (2019) All-trans Retinoic Acid as a Versatile Cytosolic Signal Modulator Mediated by CRABP1. Int. J. Mol. Sci. 10.3390/ijms20153610

4. Dowling, J. E. (2020) Vitamin A: its many roles-from vision and synaptic plasticity to infant mortality. J. Comp. Physiol. A Neuroethol. Sens. Neural. Behav. Physiol. 206, 389–399

5. Shete, V., and Quadro, L. (2013) Mammalian metabolism of β-carotene: gaps in knowledge. Nutrients. 5, 4849–4868

6. Kim, Y.-K., Zuccaro, M. V., Costabile, B. K., Rodas, R., and Quadro, L. (2015) Tissue- and sex-specific effects of β-carotene 15,15’ oxygenase (BCO1) on retinoid and lipid metabolism in adult and developing mice. Arch. Biochem. Biophys. 572, 11–18

7. Kedishvili, N. Y. (2016) Retinoic Acid Synthesis and Degradation. Subcell. Biochem. 81, 127–161

8. Harrison, E. H., and Kopec, R. E. (2020) Enzymology of vertebrate carotenoid oxygenases. Biochim. Biophys. Acta Mol. Cell Biol. Lipids. 1865, 158653

9. Napoli, J. L. (2020) Post-natal all-trans-retinoic acid biosynthesis. Methods Enzymol. 637, 27–54

10. Wang, C., Kane, M. A., and Napoli, J. L. (2011) Multiple retinol and retinal dehydrogenases catalyze all-trans-retinoic acid biosynthesis in astrocytes. J. Biol. Chem. 286, 6542–6553

11. Napoli, J. L. (2012) Physiological insights into all-trans-retinoic acid biosynthesis. Biochim. Biophys. Acta. 1821, 152–167

12. Krois, C. R., Vuckovic, M. G., Huang, P., Zaversnik, C., Liu, C. S., Gibson, C. E., Wheeler, M. R., Obrochta, K. M., Min, J. H., Herber, C. B., Thompson, A. C., Shah, I. D., Gordon, S. P., Hellerstein, M. K., and Napoli, J. L. (2019) RDH1 suppresses adiposity by promoting brown adipose adaptation to fasting and re-feeding. Cell. Mol. Life Sci. CMLS. 76, 2425–2447

13. Everts, H. B., Sundberg, J. P., King, L. E., and Ong, D. E. (2007) Immunolocalization of enzymes, binding proteins, and receptors sufficient for retinoic acid synthesis and signaling during the hair cycle. J. Invest. Dermatol. 127, 1593–1604

14. Everts, H. B., and Akuailou, E.-N. (2021) Retinoids in Cutaneous Squamous Cell Carcinoma. Nutrients. 10.3390/nu13010153

15. Siegenthaler, J. A., Ashique, A. M., Zarbalis, K., Patterson, K. P., Hecht, J. H., Kane, M. A., Folias, A. E., Choe, Y., May, S. R., Kume, T., Napoli, J. L., Peterson, A. S., and Pleasure, S. J. (2009) Retinoic acid from the meninges regulates cortical neuron generation. Cell. 139, 597–609

16. Rhinn, M., Schuhbaur, B., Niederreither, K., and Dollé, P. (2011) Involvement of retinol dehydrogenase 10 in embryonic patterning and rescue of its loss of function by maternal retinaldehyde treatment. Proc. Natl. Acad. Sci. U. S. A. 108, 16687–16692

17. Bonney, S., Harrison-Uy, S., Mishra, S., MacPherson, A. M., Choe, Y., Li, D., Jaminet, S.-C., Fruttiger, M., Pleasure, S. J., and Siegenthaler, J. A. (2016) Diverse Functions of Retinoic Acid in Brain Vascular Development. J. Neurosci. Off. J. Soc. Neurosci. 36, 7786–7801

18. Tong, M.-H., Yang, Q.-E., Davis, J. C., and Griswold, M. D. (2013) Retinol dehydrogenase 10 is indispensible for spermatogenesis in juvenile males. Proc. Natl. Acad. Sci. U. S. A. 110, 543–548

19. Wang, S., Yu, J., Kane, M. A., and Moise, A. R. (2020) Modulation of retinoid signaling: therapeutic opportunities in organ fibrosis and repair. Pharmacol. Ther. 205, 107415

20. Yang, D., Vuckovic, M. G., Smullin, C. P., Kim, M., Lo, C. P.-S., Devericks, E., Yoo, H. S., Tintcheva, M., Deng, Y., and Napoli, J. L. (2018) Modest Decreases in Endogenous All-trans-Retinoic Acid Produced by a Mouse Rdh10 Heterozygote Provoke Major Abnormalities in Adipogenesis and Lipid Metabolism. Diabetes. 67, 662–673

21. Wang, S., Yu, J., Jones, J. W., Pierzchalski, K., Kane, M. A., Trainor, P. A., Xavier-Neto, J., and Moise, A. R. (2018) Retinoic acid signaling promotes the cytoskeletal rearrangement of embryonic epicardial cells. FASEB J. Off. Publ. Fed. Am. Soc. Exp. Biol. 32, 3765–3781

22. Wang, S., and Moise, A. R. (2019) Recent insights on the role and regulation of retinoic acid signaling during epicardial development. Genes. N. Y. N 2000. 57, e23303

23. Loiselle, D. S., Johnston, C. M., Han, J.-C., Nielsen, P. M. F., and Taberner, A. J. (2016) Muscle heat: a window into the thermodynamics of a molecular machine. Am. J. Physiol. Heart Circ. Physiol. 310, H311–325

24. Petersen, M. C., and Shulman, G. I. (2018) Mechanisms of Insulin Action and Insulin Resistance. Physiol. Rev. 98, 2133–2223

25. Laurens, C., de Glisezinski, I., Larrouy, D., Harant, I., and Moro, C. (2020) Influence of Acute and Chronic Exercise on Abdominal Fat Lipolysis: An Update. Front. Physiol. 11, 575363

26. Froeschlé, A., Alric, S., Kitzmann, M., Carnac, G., Auradé, F., Rochette-Egly, C., and Bonnieu, A. (1998) Retinoic acid receptors and muscle b-HLH proteins: partners in retinoid-induced myogenesis. Oncogene. 16, 3369–3378

27. Hamade, A., Deries, M., Begemann, G., Bally-Cuif, L., Genêt, C., Sabatier, F., Bonnieu, A., and Cousin, X. (2006) Retinoic acid activates myogenesis in vivo through Fgf8 signalling. Dev. Biol. 289, 127–140

28. Kennedy, K. A. M., Porter, T., Mehta, V., Ryan, S. D., Price, F., Peshdary, V., Karamboulas, C., Savage, J., Drysdale, T. A., Li, S.-C., Bennett, S. A. L., and Skerjanc, I. S. (2009) Retinoic acid enhances skeletal muscle progenitor formation and bypasses inhibition by bone morphogenetic protein 4 but not dominant negative beta-catenin. BMC Biol. 7, 67

29. Lamarche, É., Lala-Tabbert, N., Gunanayagam, A., St-Louis, C., and Wiper-Bergeron, N. (2015) Retinoic acid promotes myogenesis in myoblasts by antagonizing transforming growth factor-beta signaling via C/EBPβ. Skelet. Muscle. 5, 8

30. El Haddad, M., Jean, E., Turki, A., Hugon, G., Vernus, B., Bonnieu, A., Passerieux, E., Hamade, A., Mercier, J., Laoudj-Chenivesse, D., and Carnac, G. (2012) Glutathione peroxidase 3, a new retinoid target gene, is crucial for human skeletal muscle precursor cell survival. J. Cell Sci. 125, 6147–6156

31. Ryan, T., Liu, J., Chu, A., Wang, L., Blais, A., and Skerjanc, I. S. (2012) Retinoic acid enhances skeletal myogenesis in human embryonic stem cells by expanding the premyogenic progenitor population. Stem Cell Rev. Rep. 8, 482–493

32. Di Rocco, A., Uchibe, K., Larmour, C., Berger, R., Liu, M., Barton, E. R., and Iwamoto, M. (2015) Selective Retinoic Acid Receptor γ Agonists Promote Repair of Injured Skeletal Muscle in Mouse. Am. J. Pathol. 185, 2495–2504

33. Li, X.-H., Kakkad, B., and Ong, D. E. (2004) Estrogen directly induces expression of retinoic acid biosynthetic enzymes, compartmentalized between the epithelium and underlying stromal cells in rat uterus. Endocrinology. 145, 4756–4762

34. Mehlem, A., Hagberg, C. E., Muhl, L., Eriksson, U., and Falkevall, A. (2013) Imaging of neutral lipids by oil red O for analyzing the metabolic status in health and disease. Nat. Protoc. 8, 1149–1154

35. Zhao, Y.-X., Pan, J.-B., Wang, Y.-N., Zou, Y., Guo, L., Tang, Q.-Q., and Qian, S.-W. (2019) Stimulation of histamine H4 receptor participates in cold-induced browning of subcutaneous white adipose tissue. Am. J. Physiol. Endocrinol. Metab. 317, E1158–E1171

36. Kane, M. A., Folias, A. E., Wang, C., and Napoli, J. L. (2008) Quantitative profiling of endogenous retinoic acid in vivo and in vitro by tandem mass spectrometry. Anal. Chem. 80, 1702–1708

37. Arnold, S. L. M., Amory, J. K., Walsh, T. J., and Isoherranen, N. (2012) A sensitive and specific method for measurement of multiple retinoids in human serum with UHPLC-MS/MS. J. Lipid Res. 53, 587–598

38. Clarke, K., Ricciardi, S., Pearson, T., Bharudin, I., Davidsen, P. K., Bonomo, M., Brina, D., Scagliola, A., Simpson, D. M., Beynon, R. J., Khanim, F., Ankers, J., Sarzynski, M. A., Ghosh, S., Pisconti, A., Rozman, J., Hrabe de Angelis, M., Bunce, C., Stewart, C., Egginton, S., Caddick, M., Jackson, M., Bouchard, C., Biffo, S., and Falciani, F. (2017) The Role of Eif6 in Skeletal Muscle Homeostasis Revealed by Endurance Training Coexpression Networks. Cell Rep. 21, 1507–1520

39. Rexer, B. N., and Ong, D. E. (2002) A novel short-chain alcohol dehydrogenase from rats with retinol dehydrogenase activity, cyclically expressed in uterine epithelium. Biol. Reprod. 67, 1555–1564

40. Markova, N. G., Pinkas-Sarafova, A., Karaman-Jurukovska, N., Jurukovski, V., and Simon, M. (2003) Expression pattern and biochemical characteristics of a major epidermal retinol dehydrogenase. Mol. Genet. Metab. 78, 119–135

41. Everts, H. B., King, L. E., Sundberg, J. P., and Ong, D. E. (2004) Hair cycle-specific immunolocalization of retinoic acid synthesizing enzymes Aldh1a2 and Aldh1a3 indicate complex regulation. J. Invest. Dermatol. 123, 258–263

42. Yoo, H. S., and Napoli, J. L. (2019) Quantification of Dehydroepiandrosterone, 17β-Estradiol, Testosterone, and Their Sulfates in Mouse Tissues by LC-MS/MS. Anal. Chem. 91, 14624–14630

43. Wood, G. A., Fata, J. E., Watson, K. L. M., and Khokha, R. (2007) Circulating hormones and estrous stage predict cellular and stromal remodeling in murine uterus. Reprod. Camb. Engl. 133, 1035–1044

44. Langley, B., Thomas, M., Bishop, A., Sharma, M., Gilmour, S., and Kambadur, R. (2002) Myostatin inhibits myoblast differentiation by down-regulating MyoD expression. J. Biol. Chem. 277, 49831–49840

45. Gomes, M. D., Lecker, S. H., Jagoe, R. T., Navon, A., and Goldberg, A. L. (2001) Atrogin-1, a muscle-specific F-box protein highly expressed during muscle atrophy. Proc. Natl. Acad. Sci. U. S. A. 98, 14440–14445

46. Gumucio, J. P., and Mendias, C. L. (2013) Atrogin-1, MuRF-1, and sarcopenia. Endocrine. 43, 12–21

47. Yaghoob Nezhad, F., Verbrugge, S. A. J., Schönfelder, M., Becker, L., Hrabě de Angelis, M., and Wackerhage, H. (2019) Genes Whose Gain or Loss-of-Function Increases Endurance Performance in Mice: A Systematic Literature Review. Front. Physiol. 10, 262

48. Schiaffino, S., and Reggiani, C. (2011) Fiber types in mammalian skeletal muscles. Physiol. Rev. 91, 1447–1531

49. Folker, E. S., and Baylies, M. K. (2013) Nuclear positioning in muscle development and disease. Front. Physiol. 4, 363

50. Brina, D., Miluzio, A., Ricciardi, S., Clarke, K., Davidsen, P. K., Viero, G., Tebaldi, T., Offenhäuser, N., Rozman, J., Rathkolb, B., Neschen, S., Klingenspor, M., Wolf, E., Gailus-Durner, V., Fuchs, H., Hrabe de Angelis, M., Quattrone, A., Falciani, F., and Biffo, S. (2015) eIF6 coordinates insulin sensitivity and lipid metabolism by coupling translation to transcription. Nat. Commun. 6, 8261

51. Zurlo, F., Larson, K., Bogardus, C., and Ravussin, E. (1990) Skeletal muscle metabolism is a major determinant of resting energy expenditure. J. Clin. Invest. 86, 1423–1427

52. Gaemers, I. C., Van Pelt, A. M., Themmen, A. P., and De Rooij, D. G. (1998) Isolation and characterization of all-trans-retinoic acid-responsive genes in the rat testis. Mol. Reprod. Dev. 50, 1–6

53. Watson, J., Lee, M., and Garcia-Casal, M. N. (2018) Consequences of Inadequate Intakes of Vitamin A, Vitamin B12, Vitamin D, Calcium, Iron, and Folate in Older Persons. Curr. Geriatr. Rep. 7, 103–113

54. Rousseau, C., Nichol, J. N., Pettersson, F., Couture, M.-C., and Miller, W. H. (2004) ERbeta sensitizes breast cancer cells to retinoic acid: evidence of transcriptional crosstalk. Mol. Cancer Res. MCR. 2, 523–531

55. Miro Estruch, I., de Haan, L. H. J., Melchers, D., Houtman, R., Louisse, J., Groten, J. P., and Rietjens, I. M. C. M. (2018) The effects of all-trans retinoic acid on estrogen receptor signaling in the estrogen-sensitive MCF/BUS subline. J. Recept. Signal Transduct. Res. 38, 112–121

56. Ombra, M. N., Di Santi, A., Abbondanza, C., Migliaccio, A., Avvedimento, E. V., and Perillo, B. (2013) Retinoic acid impairs estrogen signaling in breast cancer cells by interfering with activation of LSD1 via PKA. Biochim. Biophys. Acta. 1829, 480–486

57. Nakajima, T., Sato, T., Iguchi, T., and Takasugi, N. (2019) Retinoic acid signaling determines the fate of the uterus from the mouse Müllerian duct. Reprod. Toxicol. Elmsford N. 86, 56–61

58. Hevener, A. L., Zhou, Z., Moore, T. M., Drew, B. G., and Ribas, V. (2018) The impact of ERα action on muscle metabolism and insulin sensitivity - Strong enough for a man, made for a woman. Mol. Metab. 15, 20–34

59. Huang, H. F., Li, M. T., Von Hagen, S., Zhang, Y. F., and Irwin, R. J. (1997) Androgen modulation of the messenger ribonucleic acid of retinoic acid receptors in the prostate, seminal vesicles, and kidney in the rat. Endocrinology. 138, 553–559

60. Zhuang, Y. H., Bläuer, M., Ylikomi, T., and Tuohimaa, P. (1997) Spermatogenesis in the vitamin A-deficient rat: possible interplay between retinoic acid receptors, androgen receptor and inhibin alpha-subunit. J. Steroid Biochem. Mol. Biol. 60, 67–76

61. Buckingham, M., and Rigby, P. W. J. (2014) Gene regulatory networks and transcriptional mechanisms that control myogenesis. Dev. Cell. 28, 225–238

62. Flynn, J. M., Meadows, E., Fiorotto, M., and Klein, W. H. (2010) Myogenin regulates exercise capacity and skeletal muscle metabolism in the adult mouse. PloS One. 5, e13535

63. Maden, M. (2020) RA Signaling in Limb Development and Regeneration in Different Species. Subcell. Biochem. 95, 87–117

64. Edwards, M. K., and McBurney, M. W. (1983) The concentration of retinoic acid determines the differentiated cell types formed by a teratocarcinoma cell line. Dev. Biol. 98, 187–191

65. Langille, R. M., Paulsen, D. F., and Solursh, M. (1989) Differential effects of physiological concentrations of retinoic acid in vitro on chondrogenesis and myogenesis in chick craniofacial mesenchyme. Differ. Res. Biol. Divers. 40, 84–92

66. Momoi, T., Miyagawa-Tomita, S., Nakamura, S., Kimura, I., and Momoi, M. (1992) Retinoic acid ambivalently regulates the expression of MyoD1 in the myogenic cells in the limb buds of the early developmental stages. Biochem. Biophys. Res. Commun. 187, 245–253

67. Olson, J. M., Ameer, M. A., and Goyal, A. (2020) Vitamin A Toxicity. in StatPearls, StatPearls Publishing, Treasure Island (FL), [online] http://www.ncbi.nlm.nih.gov/books/NBK532916/ (Accessed August 25, 2020)

68. Obrochta, K. M., Kane, M. A., and Napoli, J. L. (2014) Effects of diet and strain on mouse serum and tissue retinoid concentrations. PloS One. 9, e99435

69. Chen, J., and Li, Q. (2016) Implication of retinoic acid receptor selective signaling in myogenic differentiation. Sci. Rep. 6, 18856

70. Manor, D., Shmidt, E. N., Budhu, A., Flesken-Nikitin, A., Zgola, M., Page, R., Nikitin, A. Y., and Noy, N. (2003) Mammary carcinoma suppression by cellular retinoic acid binding protein-II. Cancer Res. 63, 4426–4433

